# Autism-associated protein POGZ maintains embryonic stem cells by association with esBAF and HP1γ

**DOI:** 10.1101/2021.02.07.430173

**Authors:** Xiaoyun Sun, Linxi Cheng, Yuhua Sun

## Abstract

*POGZ*, which encodes a multi-domain transcription factor, has been found frequently mutated in neurodevelopmental disorders, particularly autism spectrum disorder (ASD) and intellectual disability (ID). However, little is known about its functions in embryonic stem cells (ESCs) and in transcriptional regulation. Here, we show that POGZ plays key roles in the maintenance of ESCs by association with the SWI-SNF (BAF) chromatin remodeler complex and heterochromatin protein 1 (HP1) proteins. Loss of POGZ induces differentiation of ESCs, likely by up-regulation of primitive endoderm and mesoderm lineage genes and by down-regulation of pluripotency-related and cell cycle genes. Genome-wide binding analysis shows that POGZ is primarily localized to gene promoter and enhancer regions where POGZ is required to maintain an open chromatin. Regulation of chromatin under control of POGZ depends on esBAF complex. Furthermore, there is an extensive overlap of POGZ and OCT4 peaks genome-wide, and both factors interact with each other. We propose that POGZ is an important pluripotency-associated factor, and its absence causes failure to maintain a proper ESC-specific chromatin state and transcriptional circuitry, which eventually leads to loss of ESC phenotype. Our work provides important insights into the roles of POGZ in the maintenance of ESC identity as well as regulation of transcription, which will be useful for understanding the etiology of neurodevelopmental disorders by *POGZ* mutation.

## Introduction

Autism risk gene *POGZ* encodes a transcription factor that contains multiple domains, including a zinc-finger cluster, an HP1- binding motif, a CENPB-DB domain, and a transposase-derived DDE domain. POGZ is thought to primarily function as a transcriptional repressor as it is a reader of heterochromatin-associated mark H3K9me3 (Vermeulen et al., 2010), is associated with heterochromatin proteins (HP1s) (Nozawa et al., 2010), and inhibits transcription by an in vitro luciferase assay (Suliman-Lavie et al., 2020). Loss of function *POGZ* mutations has been frequently linked with cardiac disease, schizophrenia, microcephaly, neuroectodermal-derived intellectual disability and autism spectrum disorders (Fukai et al., 2015; Reuter et al., 2020; Stessman et al., 2016; Tan et al., 2016; White et al., 2016; Ye et al., 2015; Zhao et al., 2019). Of note, *POGZ* is one of the top recurrently mutated genes in patients with NDDs, particularly ASD and ID (De Rubeis et al., 2014; Matrumura et al., 2020; Suliman-Lavie et al., 2020; Wang et al., 2016; Zhao et al., 2019). A recent study has reported that POGZ plays an important role in hematopoiesis (Gudmundsdottir et al., 2018).

Mouse *Pogz* mRNA is abundantly expressed during early gestation and reaches its maximum expression on E9.5. *Pogz-/-* mice are early embryonic lethal, and rarely survived beyond embryonic day 16.5 (E16.5) when neurogenesis takes place (Gudmundsdottir et al., 2018). These observations suggest that POGZ may play important roles in inner cell mass and embryonic stem cell maintenance, and induction of embryonic neural cell types.

The SWI/SNF (or BAF) ATP-dependent remodelers are highly evolutionarily conserved complexes, which include the two mutually exclusive ATPase subunits SMARCA4/BRG1 and SMARCA2/BRM, and core members such as SMARCC1/BAF155, ARID1A/BAF250a and SMARCD1/BAF60a (Alver et al., 2017). The BAF complex is known to play key roles in proliferation and differentiation in a number of different tissues and cell types, including embryonic stem cells (ESCs) (Ho et al., 2008; Panamarova et al., 2016). Regulation of BAF activity can be achieved via subunit composition and by interaction with tissue-specific transcription factors via its core subunits such as BRG1 (Narayanan et al., 2018; Ninkovic et al., 2013; Xiong et al., 2015). The role of esBAF complex in the maintenance of ESCs has been firmly established. Disruption of esBAF core subunits, such as BRG1, BAF60a, BAF250a and BAF155, results in ESC self-renewal and pluripotency defects (Alajem et al., 2015; Gao et al., 2008; Gao et al., 2019; Ho et al., 2008; Kidder et al., 2009; King and Klose, 2019; Lei et al., 2015; Liu et al., 2020; Schaniel et al., 2009; Takebayashi et al., 2013; Zhang et al., 2014).

In this work, we show that POGZ plays a key role in ESC self-renewal and pluripotency, by recruiting and interacting with the esBAF complex and HP1 proteins. Loss of POGZ induces differentiation of ESCs, likely by up-regulation of differentiation genes and by down-regulation of pluripotency-related and cell cycle genes. POGZ, esBAF and core pluripotency factors are co-localized genome-wide and extensively co-occupy enhancers, including the super-enhancers. POGZ-regulated local chromatin accessibility depends on esBAF/BRG1, and they control gene expression by modulating chromatin accessibility and nucleosome occupancy.

## Results

### Generation and characterization of *Pogz-/-* ESCs

To understand the function of POGZ, we generated *Pogz-/-* ESCs by CRISPR/Cas9 technology. The guide RNAs (gRNAs) were designed targeting the exon 2 of the *Pogz* gene (Figure 1A, up). We have successfully generated at least 3 mutant alleles (1 bp deletion, 7 bp insertion, and 284 bp insertion in exon 2 of the *Pogz* gene) (Figure 1A, bottom; Figure S1A). ESCs bearing mutant allele (-1 bp, Mut1) were chosen to perform the majority of the subsequent experiments, and ESCs with other mutant alleles (+7/+284 bp for Mut2/3, respectively) were used to confirm the results when it was necessary. The qRT-PCR analysis showed that *Pogz* mRNA levels were greatly reduced in mutant ESCs compared to controls (Figure 1B). Western blotting results showed that POGZ protein was hardly detectable in mutant ESCs using POGZ antibodies of different resources (Figure 1C), which suggested that *Pogz* mutant allele are functional null.

**Figure 1.**
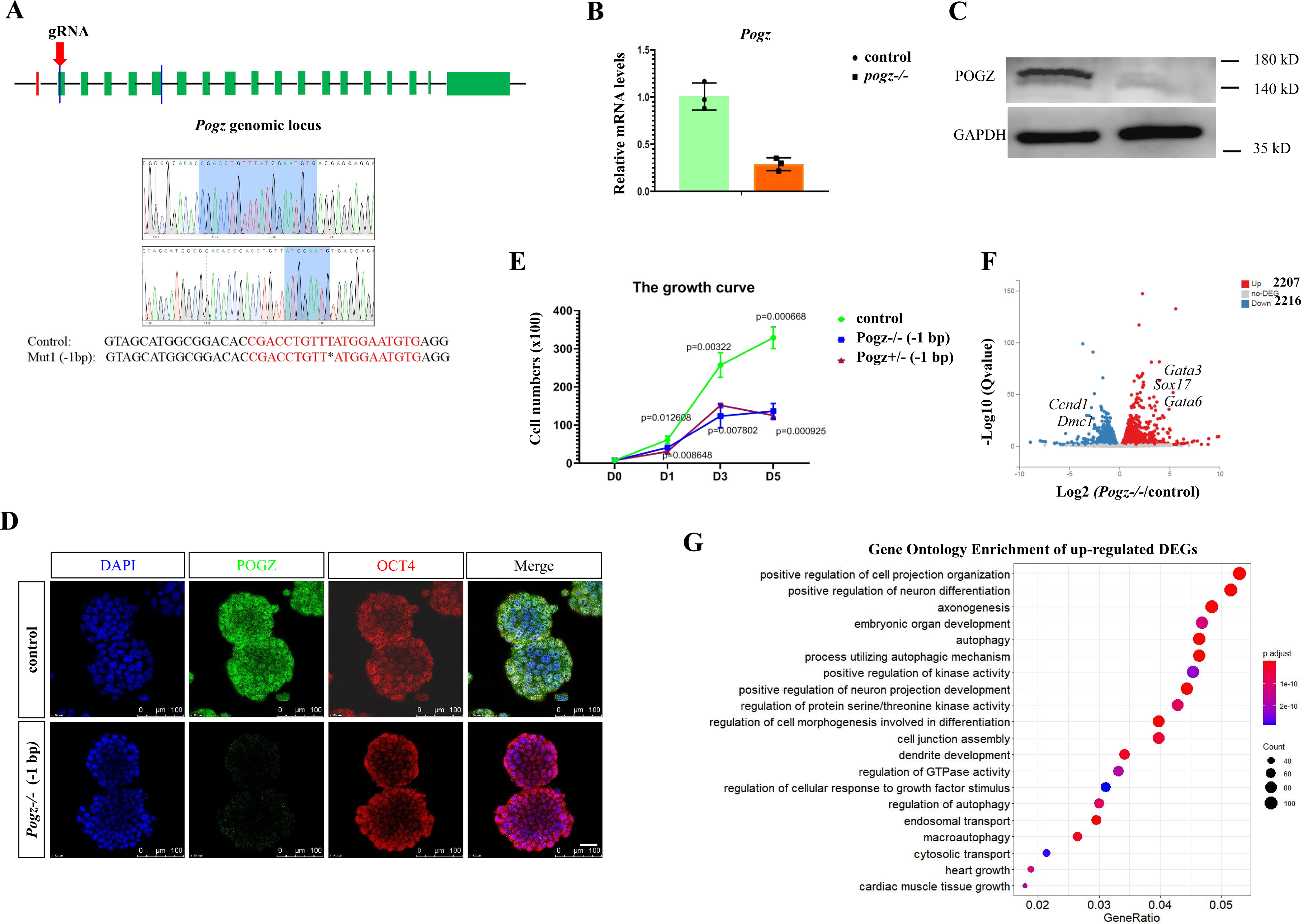
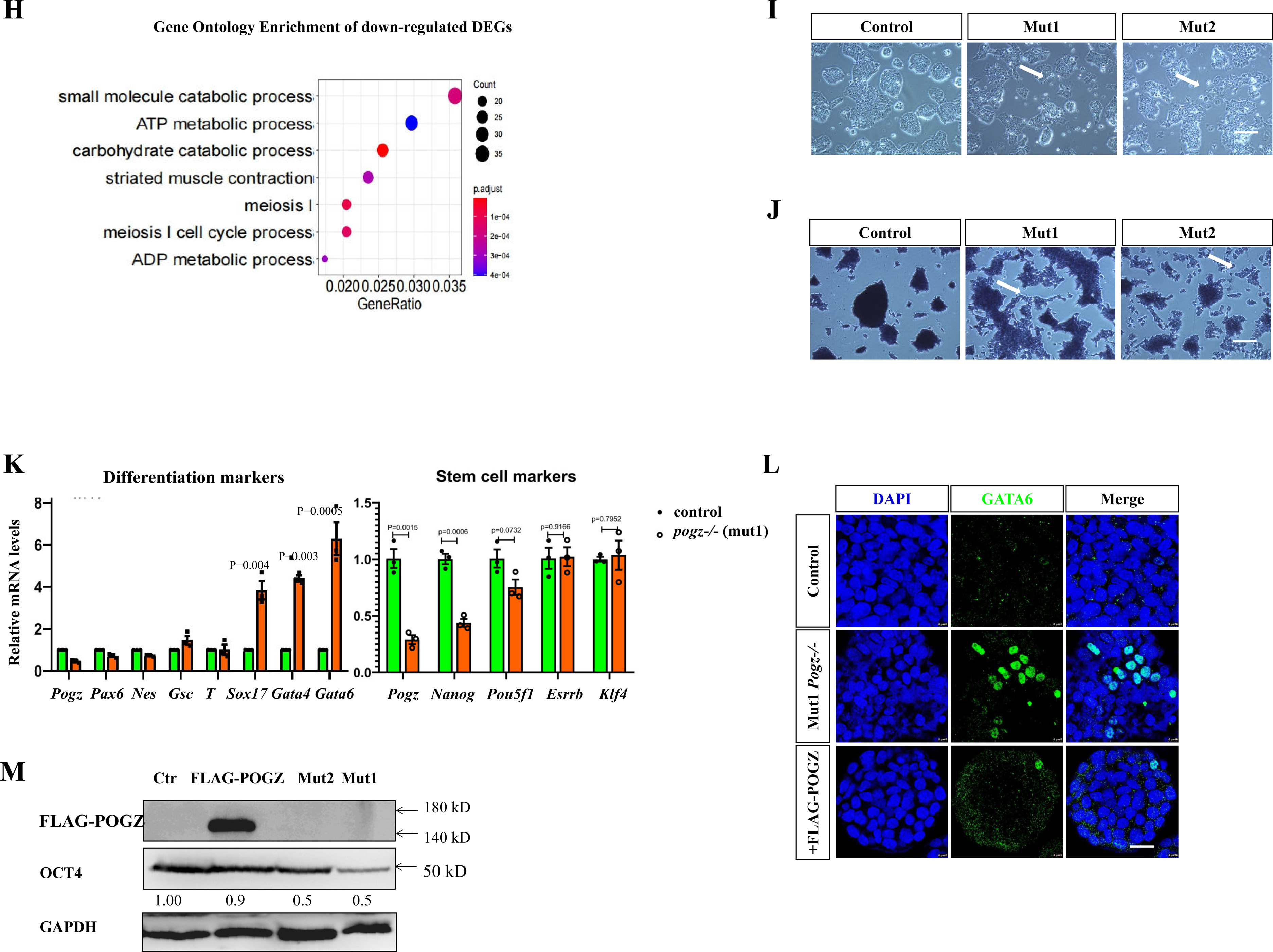
POGZ depletion leads to loss of ESC self-renewal. **A** (Up) Cartoon depicting the gRNA targeting sites at exon 2 of the mouse *Pogz* gene; (Bottom) Genotyping showing the mutant allele 1 (Mut1) of *Pogz* gene. **B** qRT-PCR showing the reduction of *Pogz* expression in mutant ESCs. **C** Western blot analysis of POGZ in control and *Pogz-/-* ESCs. **D** Double IF staining of OCT4 and POGZ showing that OCT4 levels were not significantly altered in early passage mutant ESCs (passage number is 12). Similar results were obtained for NANOG (not shown). **E** The cell growth cure showing the proliferation defects of *Pogz-/-* and *Pogz+/-* ESCs compared to control ESCs. Mut1 ESCs at passage 12 were used for the assay. **F** Valcano plot showing the up- and down-regulated genes in the control and *Pogz-/-* ESCs at passage 12. The RNA-seq experiments were repeated two times. **G** GO analysis of up-regulated DEGs between control and *Pogz-/-* ESCs. **H** GO analysis of down-regulated DEGs between control and *Pogz-/-* ESCs. **I** Representative image showing morphology of control and late passage *Pogz-/-* ESCs (passage number is 18). White arrows pointing to the flattened differentiating cells. **J** Alkaline phosphotase staining of control and *Pogz-/-* ESCs at passage 18. White arrows pointing to cells with reduced AP activities. **K** The mRNA expression levels of representative pluripotency-related, mesodermal, neuroectodermal, endodermal genes in control and *Pogz-/-* ESCs (passage number is 18). **L** IF staining of GATA6 showing that GATA6 was abnormally expressed in Mut1 *Pogz*-/- ESCs (passage number is 18). Restoring POGZ rescued the abnormal expression of Gata6. **M** WB showing that OCT4 levels were reduced in *Pogz-/-* ESCs at passage 18, which can be reversed by reintroducing FLAG-POGZ. Bar: 25 μm.

Early passage *Pogz-/-* ESCs (at passage 12) in the traditional self-renewal medium containing the LIF/KSR remained undifferentiated, as they displayed typical ESC-like domed morphology, and abundantly expressed pluripotency markers such as OCT4 (Figure 1D). In our hand, *Pogz-/-* ESCs could be maintained and expanded in undifferentiated state for at least 18 passages. However, am obvious cell proliferation defect was observed in both *Pogz-/-* and *Pogz+/-* ESCs (Figure 1E).

Immunofluorescence (IF) staining of cell division marker phospho-Histone H3 (PH3) showed that POGZ depletion decreased the number of PH3 positive cells (Figure S1B-C). Apoptosis analysis by flow cytometry revealed that POGZ depletion leads to an increase of cell death (Figure S1D). Importantly, restoring POGZ in *Pogz-/-* ESCs rescued the proliferation and apoptosis phenotype. Our results were in line with a previous report that POGZ is essential for normal mitotic progression of cells (Nozawa et al., 2010), and additionally suggested that POGZ is also required for cell survival.

To gain a global view of how POGZ is involved in the transcriptional regulation of ESCs, we performed RNA-seq analysis for control and the early passage *Pogz-/-* ESCs (Mut1 ESCs at passage 12). A total of 4423 differentially expressed genes (DEGs) were identified (Figure 1F). Of which, 2207 genes were up-regulated and 2216 genes down-regulated. Gene ontology (GO) analysis of up-regulated DEGs showed that they were enriched for terms such as regulation of nervous system development and neuronal projection, axonogenesis and muscle development (Figure 1G), and GO terms of down-regulated DEGs were enriched for metabolic process and muscle contraction (Figure 1H). KEGG analysis showed that the up-regulated DEGs were enriched for P53 and axon guidance signaling pathways, and the down-regulated DEGs were enriched for pathways in muscle contraction and metabolism (Figure S1E-F). Some pluripotency-related genes such as *Dppa3* and *Utf1* were down-regulated, however, core pluripotency genes such as *Pou5f1, Nanog, Sox2, Klf4* and *Esrrb* remained normally expressed in *Pogz-/-* ESCs, which was consistent of that early passage *Pogz-/-* ESCs could be maintained and expanded. A panel of primitive endodermal genes such as *Gata6*, *Gata4*, *Sox17*, *Sox7*, *Sparc*, *Cited*, *Cubn* and *Dab2*, mesodermal genes such as *Edn1*, *Gata3*, *Bmper*, *Nodal* and *Atf3*, and Hippo pathway genes such as *Ctgf*, *Fat1*, *Vgll3* and *Lats2*, were significantly up-regulated. By contrast, the neuroectodermal progenitor genes such as *Fgf5*, *Pax6, Olig2* and *Nes* remained largely unchanged.

Of note, positive cell cycle genes such as *E2f4*, *Ccnd1*, *Ccne1* and *Pcna*, and genes involved in segregation of chromosomes during cell division such as *Dmc1*, *Smc1* and *Sycp3*, were significantly down-regulated, while the P53 target genes such as *Cdkn2a* and *Mdm2* were up-regulated. This may explain why there are proliferation and apoptosis defects in *Pogz-/-* ESCs (Nozawa et al., 2010).

Interestingly, X chromosome-linked genes such as *Pcdh19* and *Cdkl5*, and putative imprinted genes such as *Igf2r*, *Dlk1*, *Peg3/10*, *Magel2*, *Plagl1*, *Impact*, *Sgce*, *Ndn*, *Zim1*, *Ankrd50*, *Mest* and *Zfp629* were highly represented in the up-regulated DEGs. It is known that X chromosome-linked and imprinted genes are regulated by epigenetic mechanisms such as DNA-methylation and histone modifications. This observation implied that POGZ might be involved in epigenetic regulation of X chromosome-linked and imprinted genes.

Late passaging *Pogz-/-* ESCs appeared to loss stemness progressively and finally collapsed. In our hand, the maximal passage number was 22. For instance, at passage 18, some colonies of *Pogz-/-* ESCs (Mut1/2/3) adopted a flatten morphology and lacked of tight cell contacts, indicating of spontaneous differentiation (Figure 1I). This was confirmed by alkaline phosphotase (AP) staining as the fattened cells exhibited reduced AP activities (Figure 1J). To investigate the underlying mechanism of prolonged depletion of POGZ, the expression of selected pluripotency and differentiation marker genes were examined for *Pogz-/-* ESCs at passage 18 (Mut1 and Mut2). We found that there was a decreased expression of pluripotency genes, and a significant up-regulation of primitive endoderm (PrE) markers such as *Gata6* and *Sox17* (Figure 1K; Figure S1G). We confirmed the aberrant expression of PrE marker GATA6 and the down-regulation of OCT4 at protein levels, by IF and Western blotting, respectively (Figure 1L-M; Figure S1H). Importantly, restoring FLAG-POGZ in *Pogz-/-* ESCs rescued the aberrant expression of Gata6 and OCT4 (Figure 1L-M), demonstrating the specificity of the observed phenotype.

Based on the above results, we concluded that POGZ is required for self-renew/cell cycle properties of ESCs. Its acute depletion does not lead to immediate loss of ESC phenotype, while its prolonged absence leads to ESC differentiation progressively.

### Loss of POGZ leads to compromised pluripotency

We went on to ask whether loss of POGZ leads to ESC pluripotency defects. To this end, we performed embryoid bodies (EBs) formation assay for control and early passage mutant ESCs (passage number is 12). EBs from control ESCs were round and large, whereas *Pogz-/-* ESC-derived EBs were irregular in shape and smaller in size, suggesting that there was a defect in EB formation (Figure 2A; Figure S2A).

**Figure 2.**
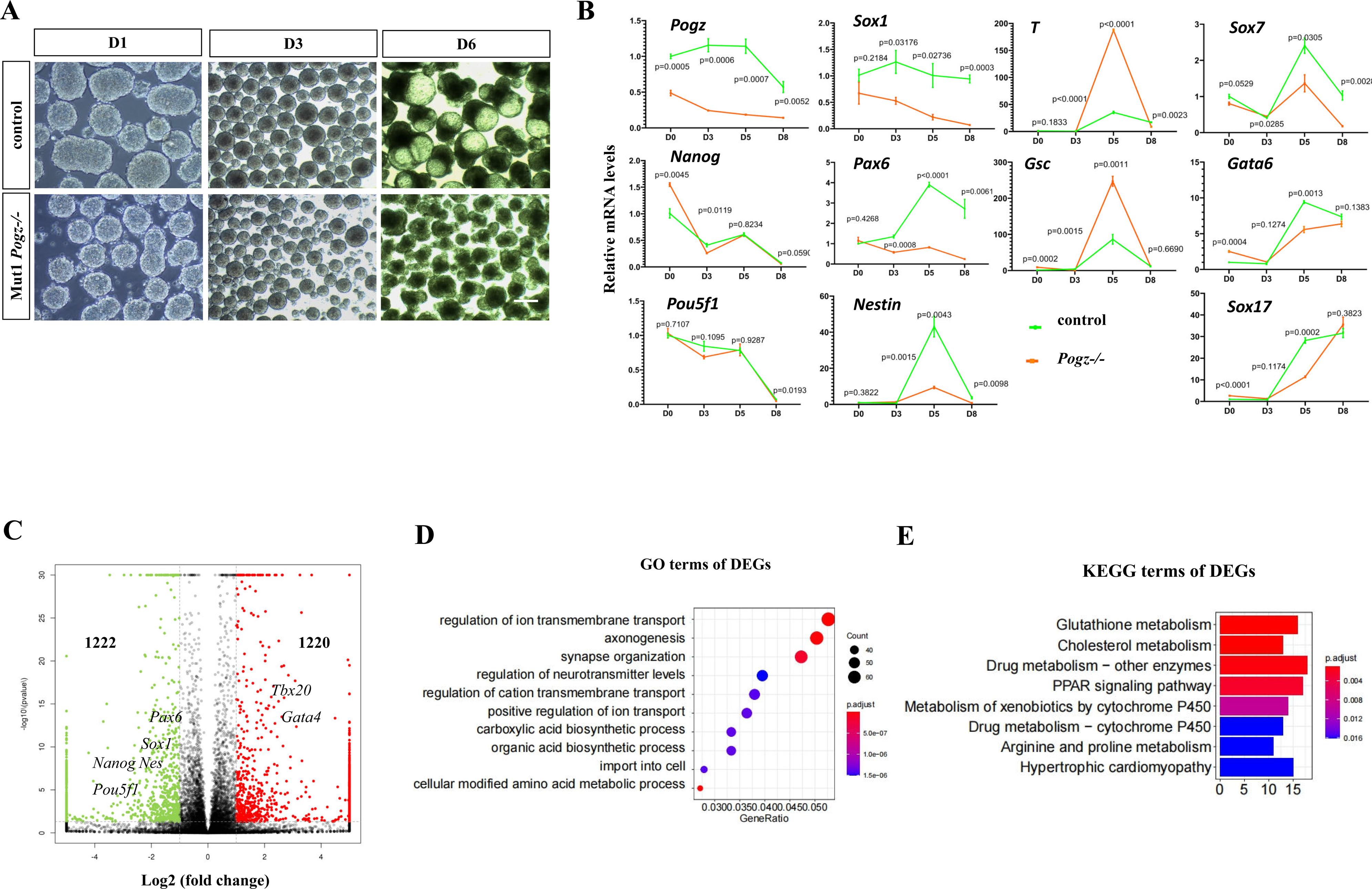
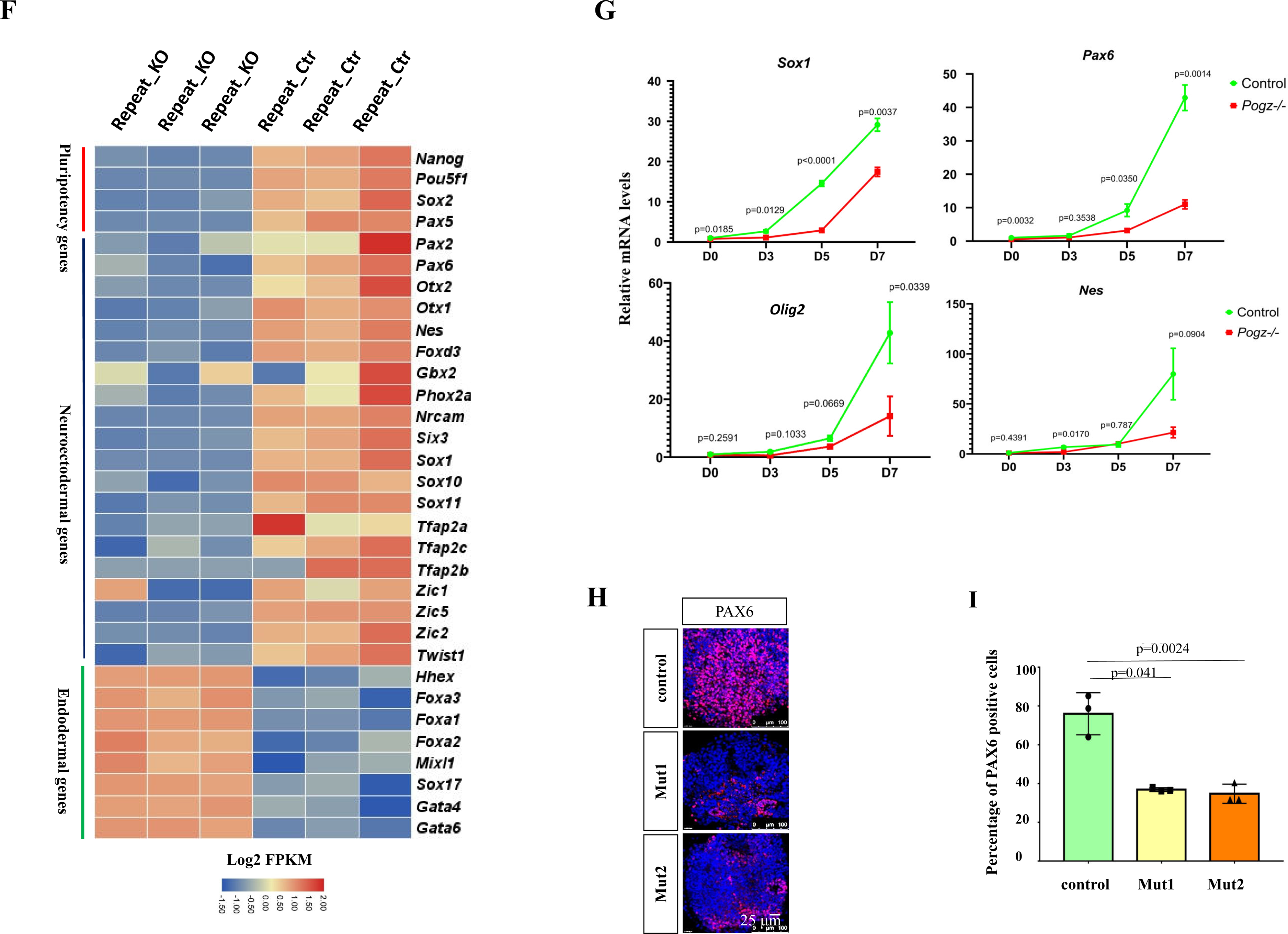
Loss of POGZ leads to compromised ESC pluripotency. **A** Morphology of day 1, 3 and 6 EBs from control and early passage Mut1 ESCs (passage is 12). **B** Time course qRT-PCR analysis of pluripotency, endodermal, mesodermal and neuroectodermal genes. **C** Valcano plot showing the up- and down-regulated genes (fold change >2) of day 6 EBs from control and *Pogz-/-* ESCs. The numbers of up- and down-regulated genes were shown as average. **D** GO analysis of DEGs of day 6 EBs from control and *Pogz-/-* ESCs (passage is 12). **E** KEGG analysis of DEGs of day 6 EBs from control and *Pogz-/-* ESCs. **F** Heat map analysis of representative endodermal, neuroectodermal and pluripotency genes. **G** Time course qRT-PCR analysis of indicated neural genes during control and Mut1 ESC directional differentiation towards neural progenitors. **H** IF results showing PAX6 positive cells in day 7 neural progenitors from control-, Mut1- and Mut2 ESCs. **I** Percentage of PAX6 positive cells in (H). Bar: 25 μm.

Next, qRT-PCR analysis was performed to carefully examine the expression of pluripotency and germ layer genes for a time course of 8 days. Compared to controls, during *Pogz-/-* ESC EB formation, neuroectodermal genes such as *Sox1*, *Nes, Pax6* and *Foxd3*, and endodermal markers such as *Gata6* and *Sox17* were markedly down-regulated (fold change> 2; *P<* 0.05) (Figure 2B; Figure S2B). By contrast, mesodermal genes such as *Tbx20*, *T* and *Gsc*, were up-regulated. Thus, the qRT-PCR analysis indicated that there was a pluripotency defect of *Pogz-/-* ESCs.

To comprehensively understand how ESC pluripotency was affected in the absence of POGZ, we performed RNA-seq analysis for day 6 control and mutant ESC-derived EBs. A total of 2,442 DEGs were identified (Figure 2C). Of which, 1,222 genes were down-regulated, and 1,220 genes were up-regulated. GO analysis of DEGs showed that they were enriched for terms such regulation of neurotransmitter levels, axon finding, synapse and neuron projection (Figure 2D). KEGG analysis of DEGs revealed the enriched terms such as pathways in regulation of cardiac development and metabolism (Figure 2E). Specifically, mesodermal genes such as *Tbx2*, *Hand2*, *Foxf1*, *Isl1, Gata4, Bmper*, *Isl1* and *Nkx2.5*, were significantly up-regulated. By contrast, neuroectodermal genes such as *Pax6*, *Nes*, *Sox1* and *Fgf5*, and pluripotency-related genes such as *Nanog*, *Sox2, Pou5f1* and *Utf1* were markedly down-regulated. The heat map analysis was used to show the representative genes (Figure 2F).

The down-regulation of neuroectodermal lineage genes during ESC EB formation was of great interest to us, considering that mutations of *POGZ* are frequently linked with neurodevelopmental disorders such as autism syndrome disorders (ASD) (Matrumura et al., 2020; Nozawa et al., 2018), which strongly suggested that POGZ is required for early neural induction by promoting neural progenitor genes (NPGs). To investigate this, we directly differentiated control and *Pogz-/-* ESCs into neural spheres for a time course of 7 days, using our previously published method (Sun et al., 2020). As shown in Figure 2G, the expression levels of NPGs were markedly decreased in *Pogz-/-* ESC derived neural progenitors compared to controls. IF staining using PAX6 antibodies further revealed that number of PAX6 positive cells in day 6 *Pogz-/-* ESC-derived neural progenitors was markedly reduced (about half of that in control ESC-derived neural progenitors) (Figure 2H-I).

Taken together, we concluded that POGZ is required for pluripotency of ESCs. In the absence of POGZ, neural progenitor genes failed to be induced, which emphasizes a critical role of POGZ in neural induction and neuronal differentiation.

### POGZ physically associates with the esBAF complex

To understand the mechanisms by which POGZ may regulate ESC self-renewal and pluripotency, we performed IP combined with mass spectrometry assay (IP-Mass Spec) to identify its potential interacting proteins. A total of 78 proteins, including the known POGZ interacting protein HP1γ/CBX3, were identified by the IP-Mass Spec (Nozawa et al., 2010; Vermeulen et al., 2010). Interestingly, members of the esBAF complex and its associated factors, including SMARCC1/BAF155, SMARCD1/BAF60a, Nono, Sfpq, Mybbp1a, Actin and ACTG1 were highly represented (Figure 3A). BAF155, BAF60a and ACTG1 are well-known members of esBAF chromatin remodeling complex and are involved in the regulation of ESC maintenance and differentiation (Alajem et al., 2015; Kim et al., 2001; Ho et al., 2008; 2009; Panamarova et al., 2016; Ramanathan et al., 2018; Schaniel et al., 2009). Nono, Sfpq and Mybbp1a are previously known esBAF-associated proteins in ESCs (Ho et al., 2009). Thus, our IP-Mass Spec experiments strongly suggested that POGZ is a cellular protein which is closely-related to the esBAF complex.

**Figure 3.**
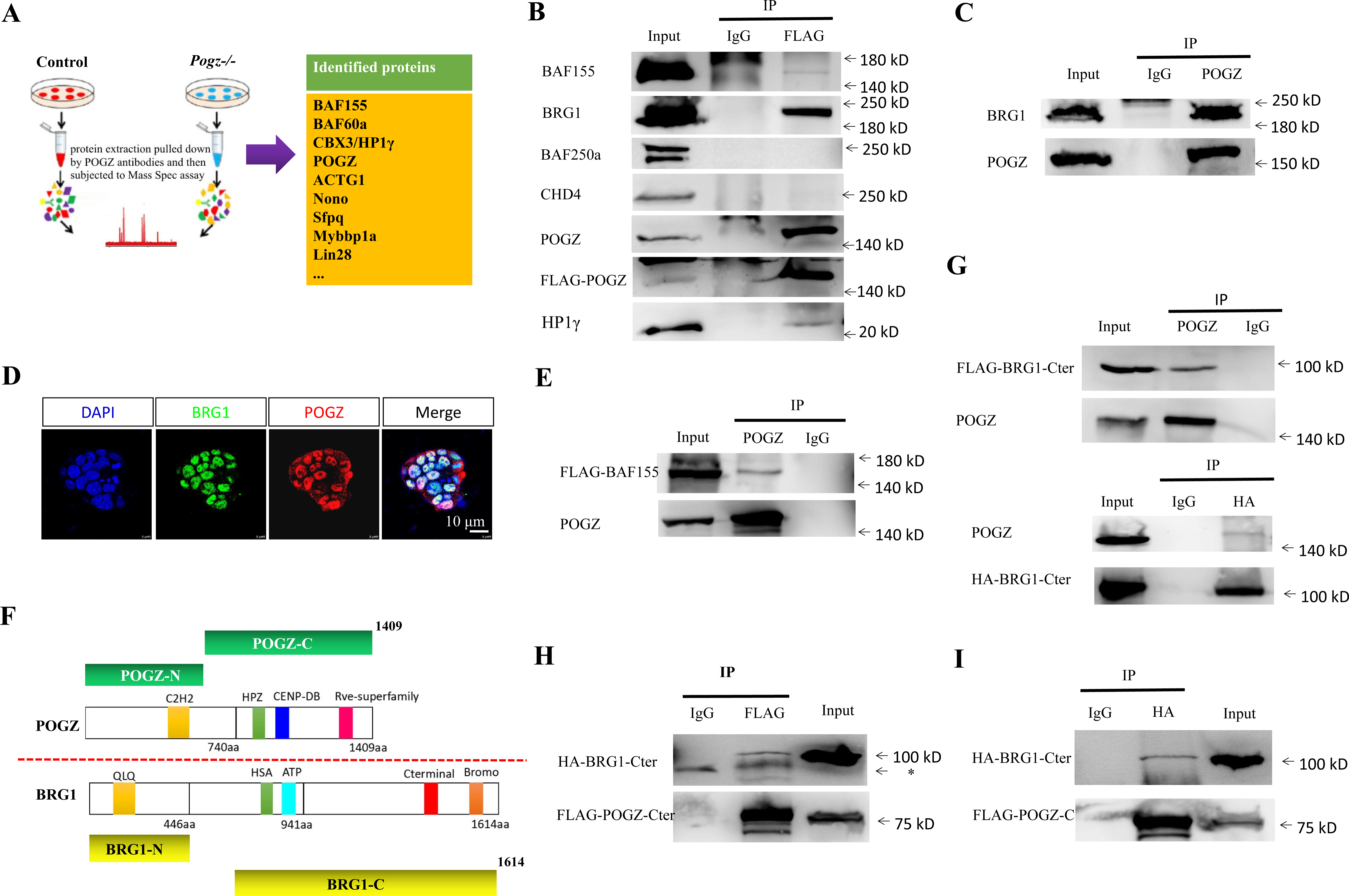
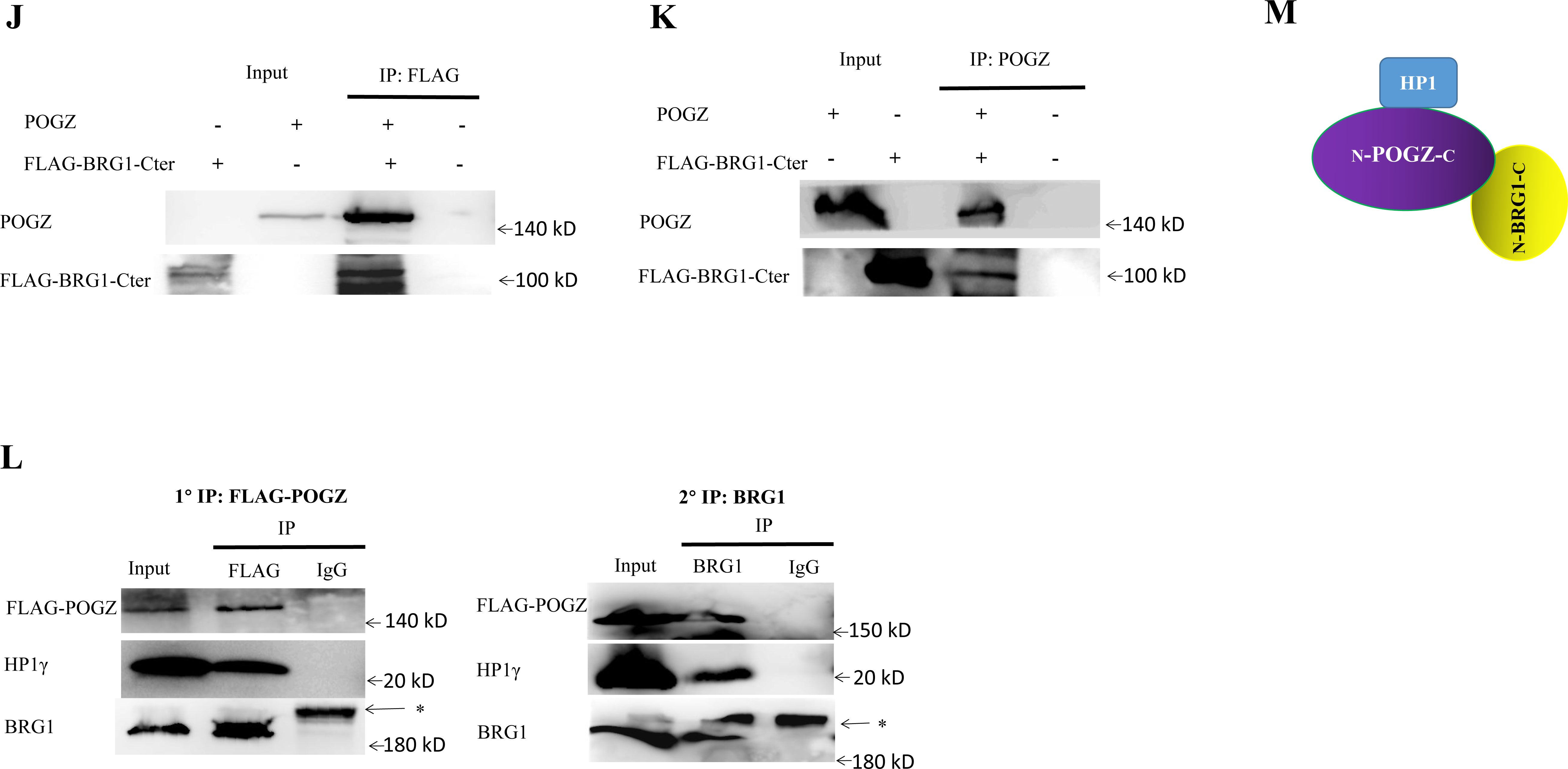
POGZ associates with esBAF and HP1. **A** Left: Cartoon showing the IP followed Mass Spec experiment. Right: the selected POGZ interacting proteins identified. IP Mass Spec experiments were repeated two times. **B** Co-IP results for interaction among FLAG-POGZ and the indicated proteins in FLAG-POGZ *Pogz-/-* ESCs. **C** Co-IP results confirming the interaction of endogenous POGZ and BRG1 in WT ESCs. **D** Double IF staining of BRG1 and POGZ in WT ESCs. **E** Co-IP results showing that POGZ interacts with FLAG-BAF155 in 293T cells. **F** Diagram showing the various truncated forms of BRG1 and POGZ used for mapping experiments. **G** Co-IP results showing that POGZ interacts with the C-terminal fragment of BRG1in 293T cells. Up: IP POGZ; WB FLAG; Bottom: IP HA; WB POGZ. **H** POGZ-Cter pulled down the C-terminal fragment of BRG1 in 293T cells. **I** The HA-BRG1-Cter pulled down FLAG-POGZ-Cter in 293T cells. **J-K** Direct interaction of in vitro synthesized POGZ and FLAG-BRG1-Cter. POGZ and FLAG-BRG1-Cter proteins were synthesized using a rabbit reticulocyte lysate system, and subjected to pull down analyses. Panel J: IP FLAG WB: POGZ; and panel K: IP POGZ WB: FLAG. **L** Sequential IP experiments showing that POGZ, BRG1 and HP1γ can for a triplex in ESCs.**M** Cartoon showing the PBH triplex. *: non-specific band. All experiments were repeated at least two times, and shown are representative images.

The IP-Mass Spec results were unexpected, as previous work have suggested that POGZ is involved in transcriptional repression and is tightly linked with repressive chromatin regulators such as CHD4 and HP1 proteins (Nozawa et al., 2010; Ostapcuk et al., 2018; Suliman-Lavie et al., 2020; Vermeulen et al., 2010). The co-IP experiments were then performed to confirm whether POGZ interacts with core members of esBAF complex as well as CHD4, using FLAG-POGZ *Pogz-/-* ESCs. The co-IP results showed that FLAG-POGZ was able to readily pull down HP1γ, which was further supported by co-localization of POGZ and HP1γ as revealed by double IF staining (Figure S3A). FLAG-POGZ appeared to strongly associate with BRG1, weakly interact with BAF155, but barely interact with BAF250a and CHD4 (Figure 3B). Furthermore, endogenous POGZ interacts with BRG1 but not CHD4 and its known binding partner ADNP (Figure 3C; Figure S3B-D). Double IF staining of BRG1 and POGZ clearly showed that they were co-localized in the nuclei of ESCs (Figure 3D). We further confirmed that POGZ interacts with FLAG-BAF155 by over-expressing them in HEK293T cells (Figure 3E). Based on the biochemistry and IF data, we concluded that POGZ is closely linked with the esBAF/BRG1 complex, but not the NuRD/CHD4 or the previously identified ChAHP members (Ostapcuk et al., 2018).

Because BRG1 is the core ATPase subunit of esBAF, we consider BRG1 to be the representative of esBAF complex in our following experiments. We went on to map the interaction domains between POGZ with BRG1. Constructs encoding the N-terminal and C-terminal fragments of POGZ and BRG1 were co-transfected into in HEK293T cells, and co-IP experiments were performed (Figure 3F). The C-terminal fragment but not the N-terminal fragment of BRG1 is responsible to interact with POGZ (Figure 3G; Figure S3E-H). Further mapping results showed that the C-terminal fragment of BRG1 mediates its interaction with POGZ-Cter but not POGZ-Nter (Figure 3H-I; Figure S3E).

Next, we asked whether POGZ physically associates with BRG1. To this end, we used the TnT in vitro translation system (Promega) to synthesize POGZ and FLAG-BRG1-Cter. When the in vitro synthesized POGZ and FLAG-BRG1-Cter were mixed together, FLAG-BRG1-Cter and POGZ can pull down each other (Figure 3K-J). We found that POGZ also directly interacts with FLAG-BAF155 using the TnT system (Figure S3I). Based on these results, we concluded that POGZ physically associates with the esBAF complex via BRG1/BAF155.

The interacting domain of POGZ (called HPZ) for HP1 proteins has been determined (Nozawa et al., 2010). We speculated that POGZ, BRG1 and HP1 may form a tripartite complex. To test this possibility, we performed the sequential immunoprecipitation experiments for POGZ, BRG1 and HP1γ. Here, HP1γ but not HP1α and β was selected because HP1γ is the only HP1 represented in our IP-Mass Spec assay and importantly, is required for proper ESC self-renewal and pluripotency similar to POGZ and BRG1 (Mattout et al., 2015; Sridharan et al., 2013; Zaidan and Sridharan, 2020). In the first round of IP, FLAG-POGZ readily pulled down BRG1. In the second round of IP, BRG1 readily pulled down HP1γ (Figure 3L). Thus, our sequential IP results strongly suggested that POGZ, BRG1 and HP1γ can form the PBH triplex (Figure 3M), although the triplex is likely a part of a much larger complexes containing other esBAF members such as BAF155 and BAF60A (for this, the PBH also stands for POGZ, esBAF and HP1).

### POGZ and core pluripotency factors are colocalized genome-wide

To understand the function of POGZ in the maintenance of ESCs and regulation of gene expression, we performed ChIP-seq experiments using different sources of commercial POGZ antibodies. Unfortunately, these commercial antibodies, although good in Western blotting and/or IF applications, failed to work in ChIP-seq experiments. We therefore performed FLAG ChIP-seq experiments using FLAG-POGZ restoring *Pogz-/-* ESC lines. The CUT&Tag, a newly developed enzyme-tethering technique that has been shown to be more efficient than the traditional ChIP-seq assay, was also utilized in parallel (Kaya-Okur et al., 2019).

A total of 7,929 peaks were identified, based on the FLAG ChIP-seq experiments. By contrast, a total of 16,728 peaks were revealed by the CUT&Tag. In addition to the larger number of POGZ peaks, the binding intensity was much stronger by CUT&Tag than by traditional ChIP-seq, which suggested that CUT&Tag outperformed the FLAG ChIP-seq. The snapshots of *Dcp1a* and *Chchd1* loci were shown for representatives (Figure S4A).

Analysis of both the CUT&Tag and FLAG ChIP-seq signals revealed that POGZ was primarily localized to the proximal Transcription Start Site (TSS) and distal intergenic regions (Figure 4A-C). At a few gene loci, POGZ peaks at distal regions were overlapped with H3K4me1 and H3K27Ac, known histone marks for poised and active enhancers, respectively (Figure S4B). This was also true at a global level (Figure 4D). In fact, we found that POGZ is highly enriched at active enhancers decorated with both H3K27ac and H3K4me1 (Figure S4C). We confirmed this by ChIP-PCR analysis of active genes such as *Pou5f1* and *Nanog* (Figure 4E).

**Figure 4.**
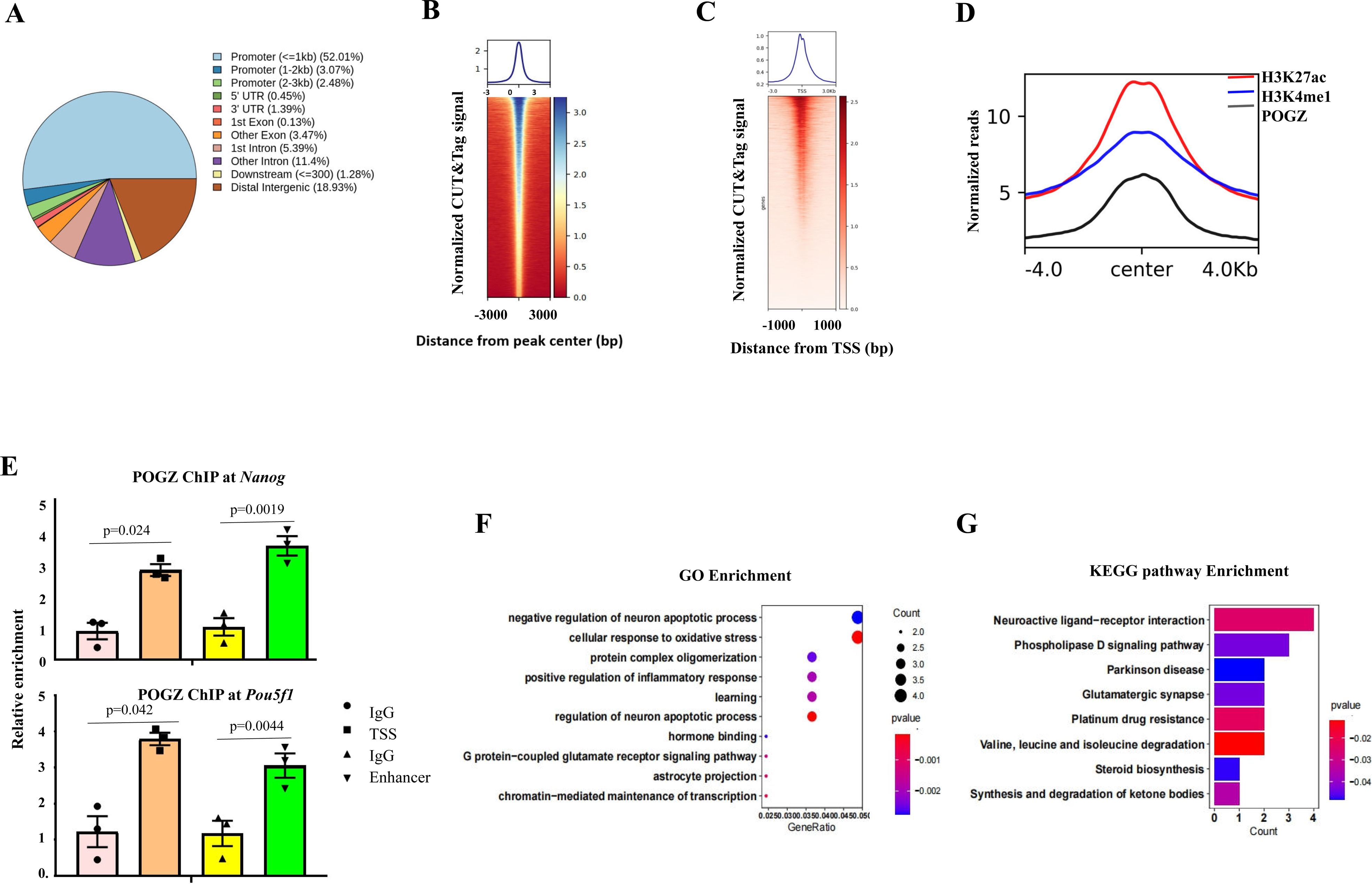
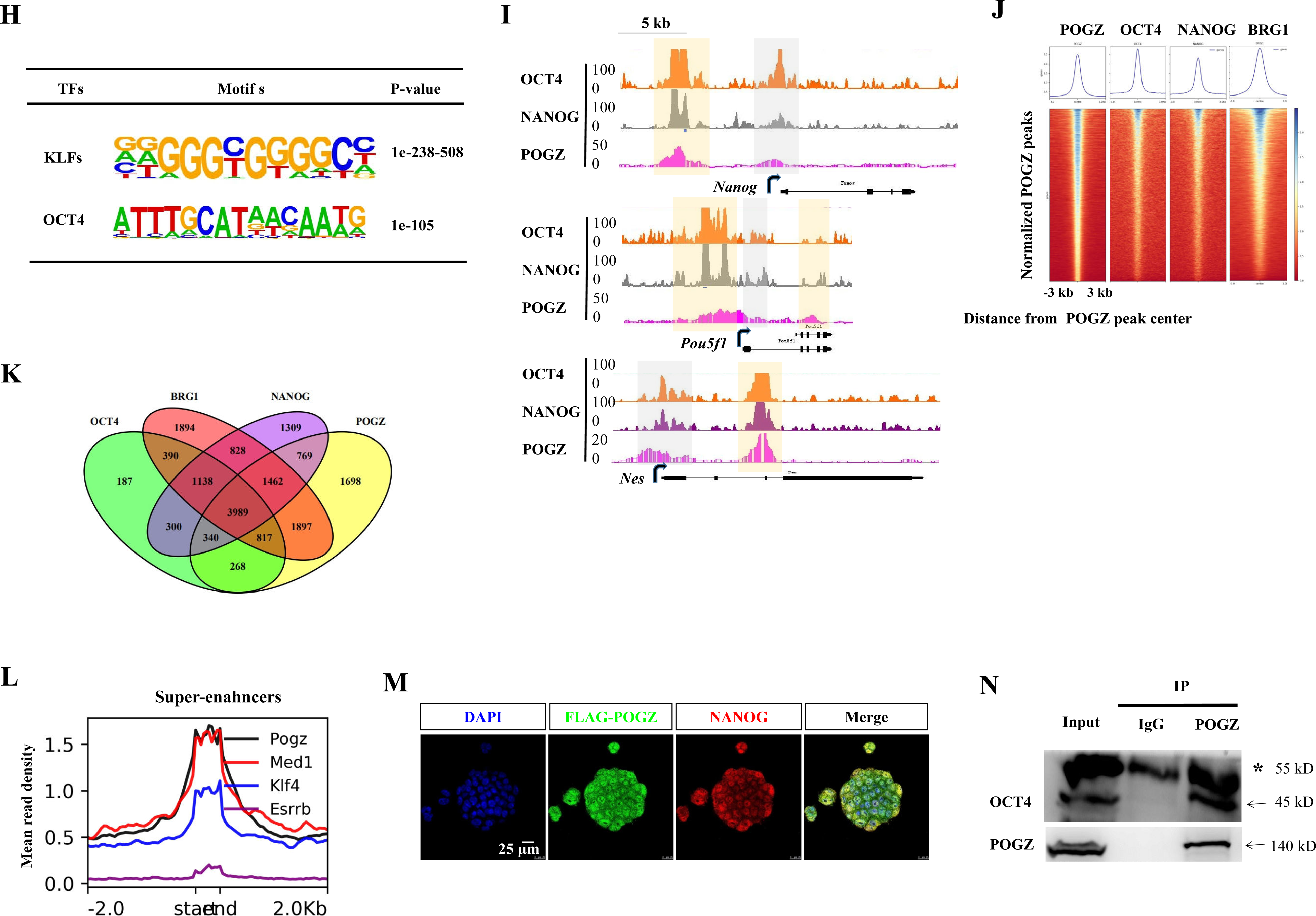
POGZ and core pluripotency factors are co-localized genome-wide. **A** Pie chart showing the distribution of POGZ binding sites genome wide, based on ChIP-seq and CUT&Tag results. **B** Heat map of POGZ CUT&Tag enrichment in a 6 kb window centered on peak midpoint of POGZ sites (n=16,728). **C** Heat map of POGZ CUT&Tag enrichment in a 6 kb window around the TSS. **D** Metaplot of POGZ, H3K4me1 and H3K27ac enrichment (normalized per million mapped reads) on ±4 kb of genes that were bound by POGZ. **E** ChIP-PCR analysis showing the enrichment of POGZ at TSS and enhancer regions at *Nanog* and *Pou5f1* loci. **F** GO analysis of POGZ-bound target genes in ESCs. **G** KEGG analysis of POGZ-bound genes in ESCs. **H** The enrichment of KLFs and OCT4/SOX2 binding motifs in POGZ-bound loci, as revealed by HOMER. **I** CUT&Tag and ChIP-seq snapshots showing the co-localization of POGZ, NANOG and OCT4 at the indicated loci. Grey: proximal TSS; Orange: distal regions. **J** Heat map of BRG1, NANOG, OCT4 and POGZ enrichment across genes bound by POGZ (n= 11000). Each row represents a 6 kb window centered on the peak midpoint. **K** A venn diagram showing the overlapping binding sites among POGZ, BRG1, NANOG and OCT4. **L** Metaplot of POGZ, MED1, ESRRB and KLF4 enrichment (normalized per million mapped reads) on ±2 kb window across all 231 super-enhancers that have been identified in ESCs by Whyte (Whyte et al. 2013). **M** Double IF staining of NANOG and FLAG-POGZ in ESCs. **N** Co-IP results showing that OCT4 interacts with POGZ in ESCs. Star pointing to the IgG heavy chain. All IF and WB experiments were repeated at least two times, and shown are representative images. Bar: 25 μm.

A total of approximately 11,000 target genes were identified for POGZ. GO analysis of POGZ-bound genes revealed the enrichment of terms such as regulation of synaptic function, regulation of neuron apoptosis, glutamate receptor activities, astrocyte projection, and regulation of chromatin (Figure 4F). KEGG analysis of POGZ target genes showed the enrichment of terms such as Parkinson diseases, glutamatergic synapse and neuroactive ligand-receptor interaction (Figure 4G). The results were in line with that POGZ is one of the most recurrently mutated genes in patients with neurodevelopmental disorders (Matrumura et al., 2020; Zhao et al., 2019).

POGZ is a typical transcription factor that is predominantly localized in nuclei, suggesting that it binds to DNA in a sequence specific manner. Motif analysis of POGZ-bound sites by HOMER revealed a series of consensus DNA motifs of known TFs (Figure 4H; Figure S4D). The top ranked motifs included SP1/2 and KLFs (Figure S4D). KLF4 in KLF family of transcription factors is a well-known pluripotency factor and is particularly enriched at super-enhancers (Whyte et al., 2013). SP1, a previously identified POGZ interactor, is involved in transcription regulation (Gunther et al., 2000). Of note, ATTTGCAT, the putative binding motif of ESC core pluripotency factors (Boyer et al., 2005; Loh et al., 2006), was enriched with high significance. It is well-known that the core pluripotency factors co-occupy both ESC-specific genes and lineage-specifying genes by binding to the ATTTGCAT motif, promoting the former and repressing the latter (Boyer et al., 2005). We found that similar to the core pluripotency factors, POGZ exhibited enrichment at ESC-specific and lineage-specifying genes, including neural progenitor genes such as *Pax6*, *Nes*, *Sox1* and *Olig2*, endodermal regulatory genes such as *Gata6* and *Sox17*, mesodermal genes such as *T* and *Gsc*, and pluripotency TF genes such as *Pou5f1* and *Nanog* (Figure 4I; Figure S4B).

As POGZ share binding motifs with the core pluripotency factors, we asked whether POGZ is co-localized with NANOG/OCT4 genome wide. To this end, we consulted the published NANOG/OCT4 ChIP-seq data sets (King and Klose, 2017). We found that there was an extensively overlap of POGZ and NANOG/OCT4 peaks genome-wide (Figure 4J-K). Interestingly, broad POGZ CUT&Tag signals (4-5 kb) were observed at distal regions of certain genes (Figure 4I; Figure S4B). This feature of binding was analogous to the previously described super-enhancers that are marked by Mediator, ESRRB, KLF4 and P300/H3K27Ac (Pott and Lieb, 2014; Whyte et al., 2013). We thus speculated that POGZ may also bind to super-enhancers. The previous study has identified 231 super-enhancers in ESCs (Whyte et al., 2013). Our global meta-analysis showed that POGZ is enriched at super-enhancers, similar to ESRRB, KLF4 and Med1 (Figure 4L). Snapshots of POGZ binding profile at selected super-enhancers were shown (Figure S4E).

The above co-localization analysis of POGZ and OCT4/NANOG revealed very high degrees of target gene overlap (Figure 4J-K), suggesting that there is a functional relationship among them. We therefore asked whether POGZ and OCT4/NANOG interact with each other in ESCs. Double IF staining showed that POGZ was clearly co-localized with NANOG in the nuclei of ESCs, similar to OCT4 (Figure 4M; Figure 1D). Importantly, the results of co-IP showed that POGZ could readily pull down endogenous OCT4 (Figure 4N).

Taken together, we concluded that POGZ is extensively co-localized with ESC master transcription factors genome-wide, and interacts with these TFs in ESCs. This raised an interesting possibility that POGZ is an important pluripotency-associated factor.

### POGZ, BRG1 and HP1 are extensively co-localized genome-wide

As POGZ interacts with esBAF and HP1 proteins in ESCs, we asked whether these factors were co-localized genome wide. We consulted the previously published ChIP-seq data for BRG1 and HP1 proteins (King and Klose, 2018; Sridharan et al., 2013). Analysis of ChIP-seq data sets showed that POGZ, BRG1 and HP1 proteins were extensively co-localized genome-wide (Figure 5A; Figure S5A). POGZ co-occupied with BRG1 and HP1 proteins at the POGZ peak center and at the proximal TSS (Figure 5B). Approximately 73% (8165/11240) of POGZ peaks were overlapped with 66% (8165/12415) of BRG1 peaks. About 59% (2510/4267) of HP1γ peaks were overlapped with 22% (2510/11240) of POGZ peaks. When examining the sites bound by all three factors, approximately 2100 peaks were identified (Figure 5C).

**Figure 5.**
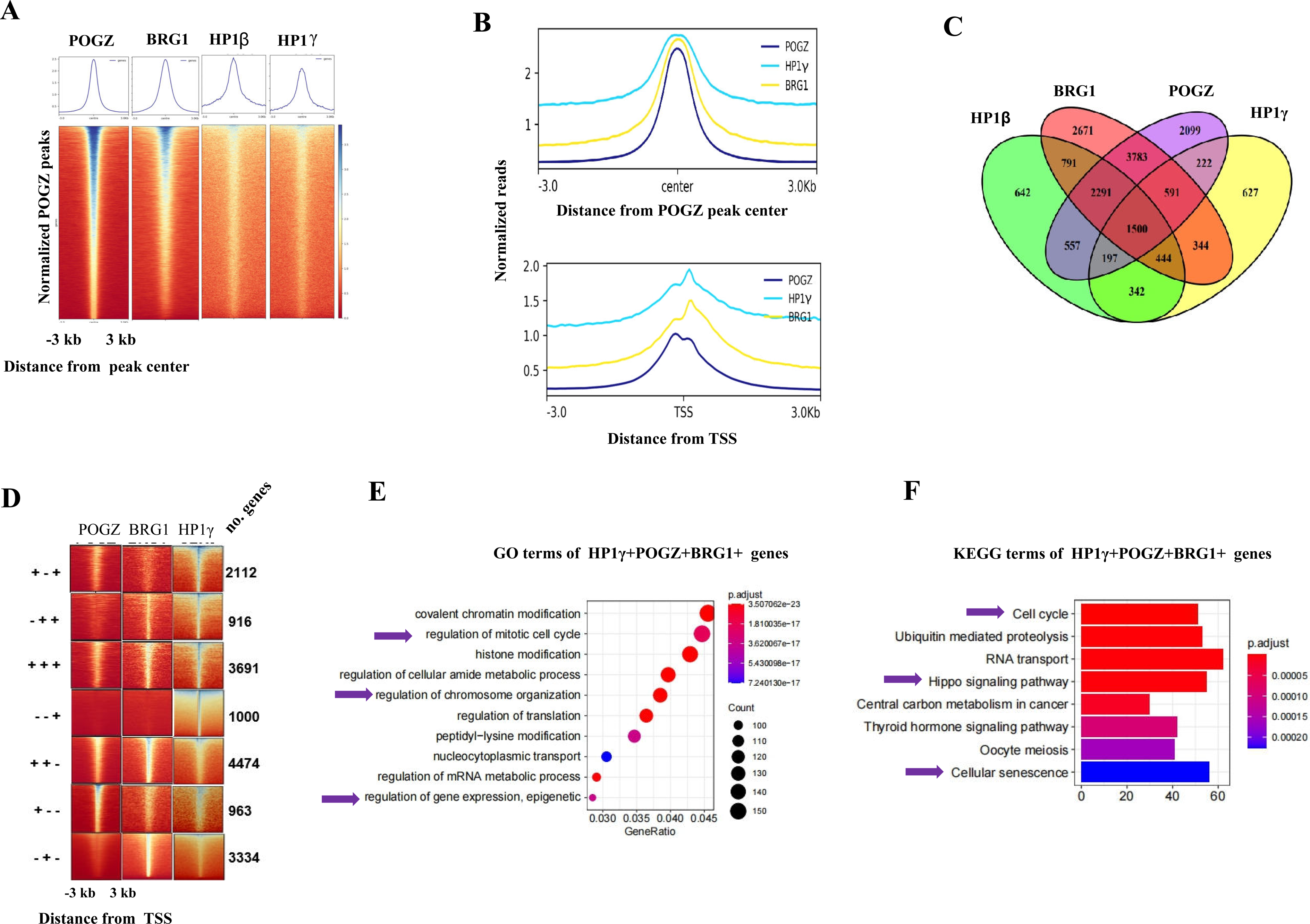
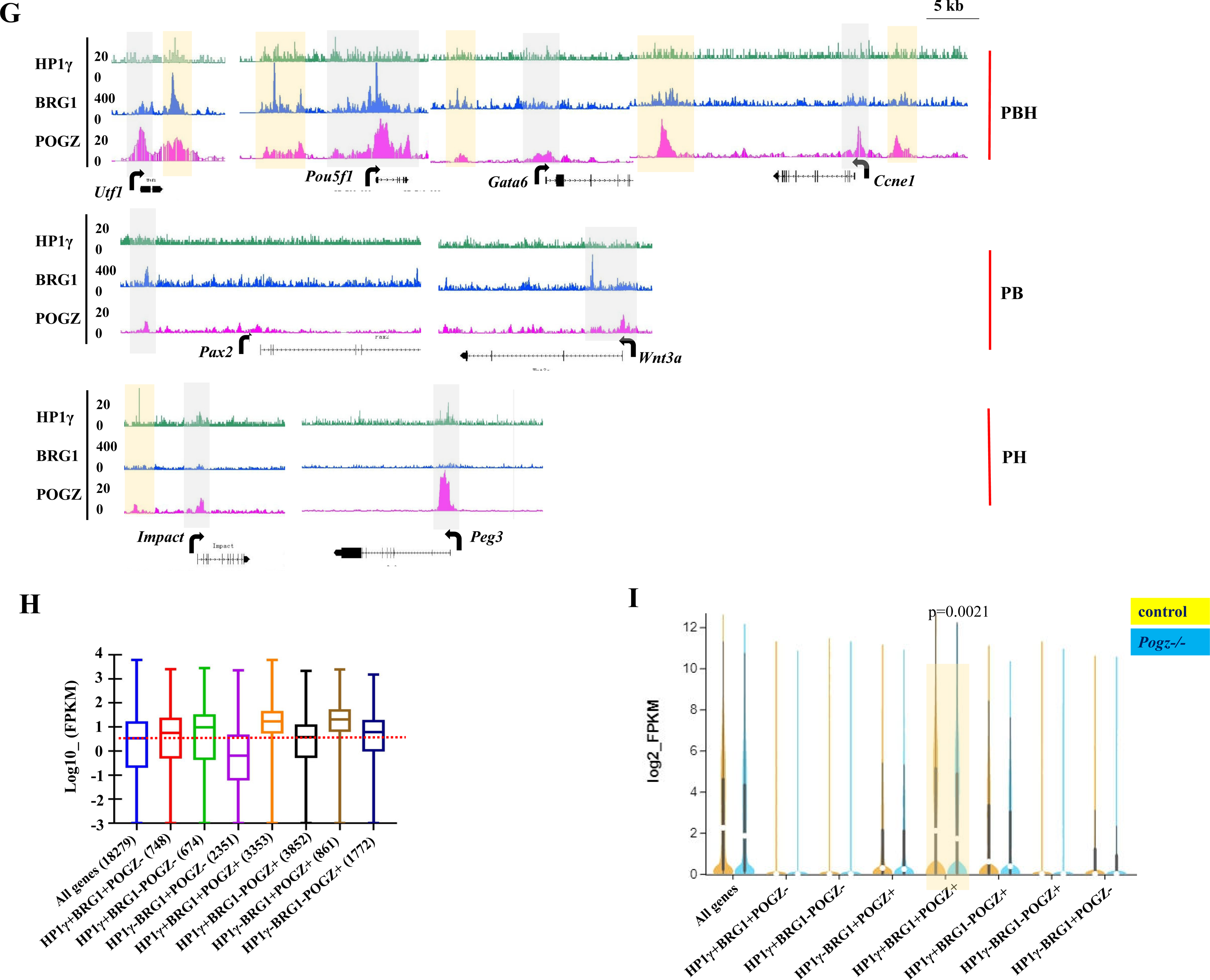
POGZ, BRG1 and HP1 are co-localized genome wide. **A** Heat map of BRG1, HP1β/CBX1, HP1γ/CBX3 and POGZ enrichment across sites bound by POGZ (n=16,728). Each row represents a 6 kb window centered on the peak midpoint. **B** (Up) Metaplot of POGZ, BRG1 and HP1γ enrichment (normalized per million mapped reads) across sites bound by POGZ; (Bottom) Metaplot of POGZ, BRG1 and HP1γ enrichment (normalized per million mapped reads) across proximal TSS. A 6 kb window centered on the peak midpoint was shown. **C** A Venn diagram showing the shared binding sites by POGZ, BRG1, HP1γ and HP1β. **D** Heat map view for distribution of POGZ, BRG1 and HP1γ/CBX3 signals in a ± 3 kb regions around the TSS. Genes therefore are classified into 7 sub-groups based on the occupancy of POGZ, BRG1 and HP1γ. **E** GO analysis of HP1γ+BRG1+POGZ+ genes. **F** KEGG analysis of HP1γ+BRG1+POGZ+ genes. **G** CUT&Tag and ChIP-seq snapshots showing the distribution of POGZ, BRG1 and HP1γ at representative PH-, PB- and PBH-bound genes. TSS (grey color) and distal enhancer regions (orange). **H** The average expression levels of the indicated gene clusters in ESCs. **I** Log2 (FPKM) showing that PBH- genes (highlighted by orange) exhibited highest expression levels in control ESCs and were mostly down-regulated among all other gene clusters in *Pogz-/-* ESCs.

Next, we plotted POGZ CUT&Tag, BRG1 and HP1γ ChIP-seq reads in a ± 3 kb region surrounding the TSS and divided POGZ-, HP1γ- and BRG1-bound genes into 7 categories (cluster A: POGZ^+^BRG1^+^HP1^+^ cluster B: POGZ^+^BRG1^-^HP1^+^, cluster C: POGZ^+^BRG1^+^HP1^-^, cluster D: POGZ^+^BRG1^-^HP1-, cluster E: POGZ^-^BRG1^+^HP1^+^, cluster F: POGZ^-^BRG1^-^HP1^+^, cluster G: POGZ^-^BRG1^+^HP1^-^) (Figure 5D). GO and KEGG analyses were performed for all gene clusters (Figure 5E-F; Figure S5B-D and not shown). For instance, KEGG revealed that cluster A genes (PBH) were enriched for terms such as cell cycle, Hippo signaling and cellular senescence, and cluster C genes (PB) were enriched for terms such as neurodegeneration and Wnt signaling pathways. Representative snapshots of cluster A/B/C genes were shown for representatives (Figure 5G).

Compared to all genes, the average mRNA expression levels of clusters A (PBH) were highest among all clusters in ESCs (Figure 5H). In fact, both HP1γ and BRG1 ChIP-seq peaks have been shown to be enriched surrounding the proximal TSS, and importantly, their enrichment levels are positively correlated with gene expression levels: namely, the higher the gene expression levels, the more enrichment of HP1γ and BRG1 at TSS (Ho et al., 2009; Zaidan and Sridharan, 2020). Thus, PBH marked highest expression genes in ESCs. Next, we analyzed the effects of loss of POGZ on expression of each cluster of genes in ESCs. We found that loss of POGZ had little effects on average expression levels of each cluster (Figure S5E). We speculated that this was likely due to the neutralization effect of up- and down-regulated genes in each cluster. We therefore examined the up-regulated and down-regulated genes, respectively, and found that cluster A genes (PBH) were significantly down-regulated in the absence of POGZ (Figure 5I). This data indicates that PBH co-occupies and facilitates a subset of highly expressing genes in ESCs.

### POGZ recruits BRG1 and HP1 to target genes

The co-localization of POGZ with BRG1 and HP1γ strongly suggested that POGZ recruits HP1γ and BRG1 to target genes. We therefore performed ChIP experiments to access the binding of HP1γ and BRG1 at POGZ-bound genes, in the presence and absence of POGZ. Pluripotency TF genes such as *Nanog* and *Pou5f1*, endodermal genes such as *Gata6*, cell cycle genes such as *Ccne1*, neuroectodermal gene*s Pax6, Nes* and *Sox1*, and Wnt and Hippo pathway genes *Myc*, *Axin2*, *Yap* and *Tead2*, were examined. The ChIP-PCR results showed that BRG1 and HP1γ levels at the majority of examined genes were significantly reduced in the absence of POGZ (Figure 6A; Figure S6A).

**Figure 6.**
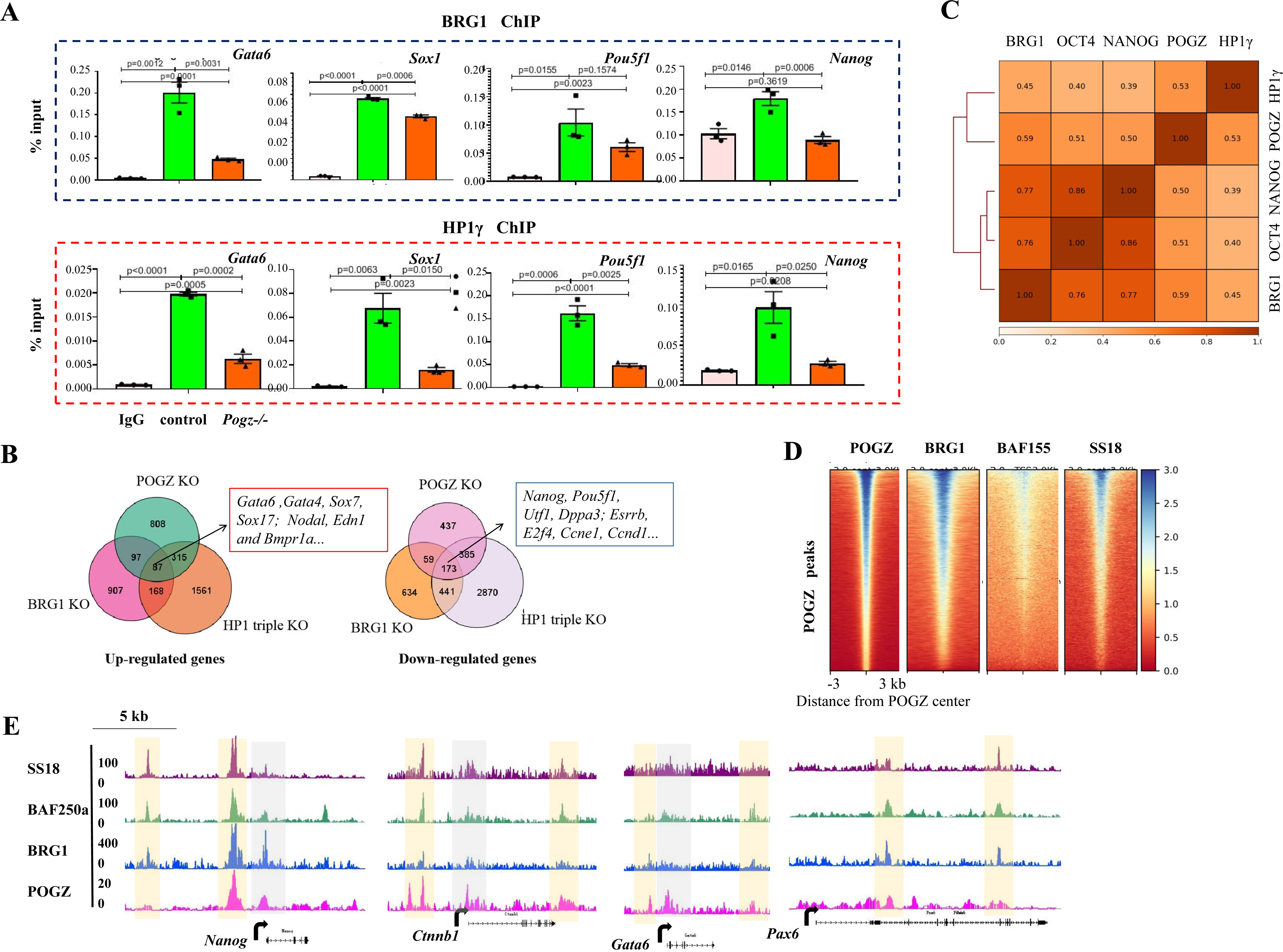
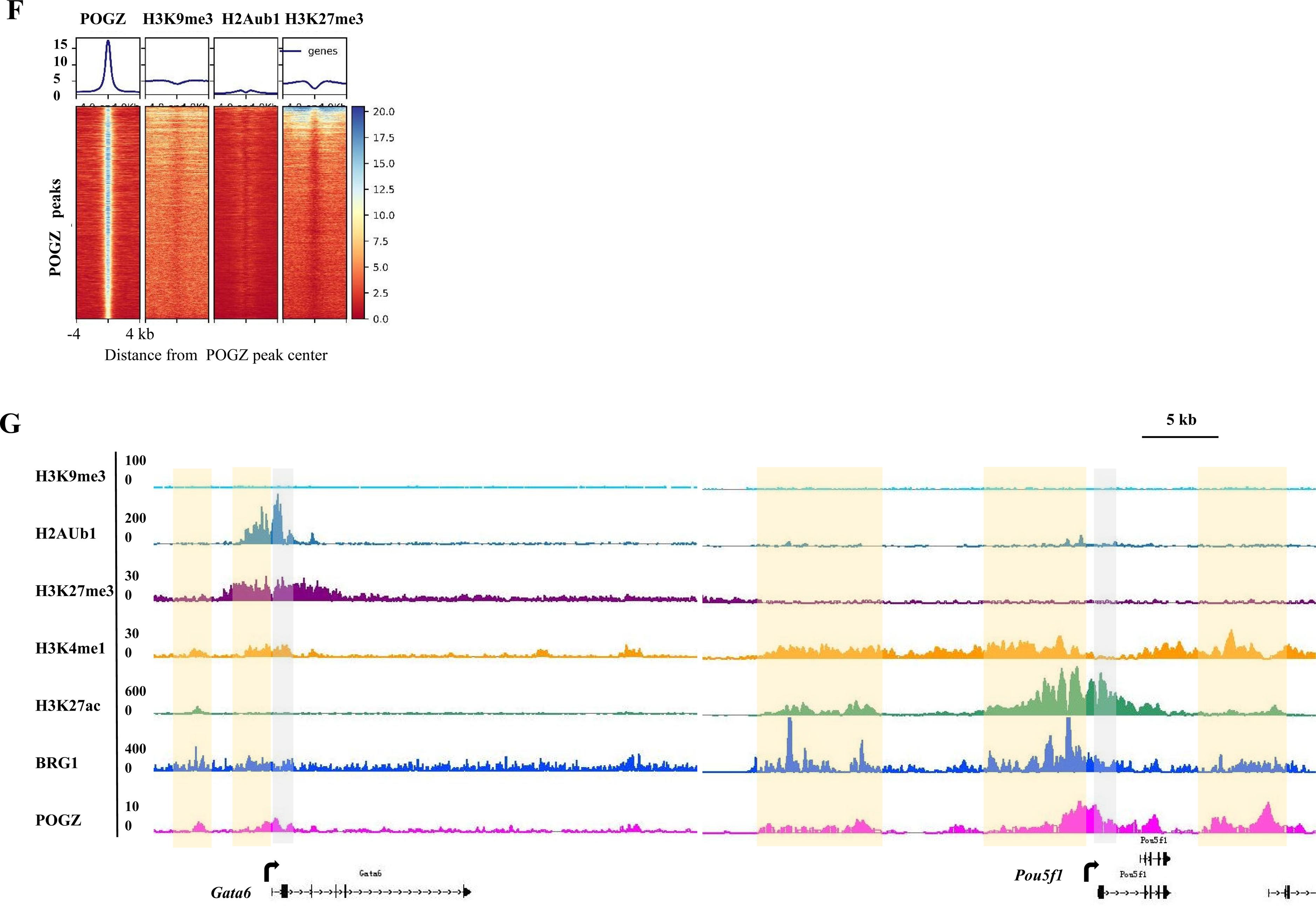
POGZ recruitment of BRG1 and HP1 is important for gene expression. **A** ChIP-PCR results showing that BRG1 and HP1γ levels were significantly reduced at the indicated genes, in early passage *Pogz-/-* ESCs (Mut1 passage 12). The up panel is for the BRG1 and bottom is for the HP1γ. **B** A Venn diagram showing the shared up-regulated and down-regulated genes (fold change> 2, and *P<* 0.05) in *Pogz-/-* (Mut1 passage 18), BRG1 depleted- and HP1 triple knockout (KO) ESCs, compared to control ESCs. Left panel: up-regulated genes; right: down-regulated genes. **C** A Pearson correlation analysis of ChIP-seq peaks revealed a high degree of co-localization of POGZ, BRG1, HP1γ, NANOG and OCT4 genome wide. **D** Heat map of POGZ, BRG1, BAF155 and SS18 enrichment across sites bound by POGZ (n=16,728). Each row represents a 6 kb window centered on the peak midpoint. **E** CUT&Tag and ChIP-seq snapshots showing the co-localization of POGZ, BRG1, BAF250a and SS18 at the indicated gene loci. Grey: proximal TSS; Orange: enhancer regions. **F** Heat map of POGZ, H3K9me3, H2Aub1 and H3K27me3 enrichment across sites bound by POGZ (n=16,728). Each row represents a 8 kb window centered on the peak midpoint. Note the depletion of H3K9me3, H2Aub1 and H3K27me3 signals at POGZ peaks. **G** CUT&Tag and ChIP-seq snapshots showing the enrichment of POGZ, BRG1, H3K9me3, H2Aub1, H3K27me3, H3K4me1 and H3K27ac at the indicated gene loci. Grey: proximal TSS; Orange: enhancer regions.

The above data predicted that PBH may support ESC stemness by controlling a cohort of common target genes. If this was the case, loss of either factor should affect the expression of similar genes or pathways. To test this hypothesis, we consulted the previously published RNA-seq data from ESCs that lack of all three isoforms of HP1 and ESCs depleted of BRG1 (Hainer et al., 2015; King and Klose, 2017; Ostapcuk et al., 2018). Bioinformatics analysis revealed a considerable shared DEGs among POGZ-, HP1- and BRG1-depleted ESCs. For instance, a total of 402 up-regulated genes and 558 down-regulated genes were shared by POGZ- and HP1-depleted ESCs, and a total of 184 up-regulated genes and 232 down-regulated genes were shared between POGZ- and BRG1-depleted ESCs (Figure 6B). When we examined the up- and down-regulated genes shared among POGZ-, BRG1- and HP1-depleted ESCs, the numbers were 87 and 173, respectively. Specifically, up-regulated genes included PrE genes such as *Gata4*, *Gata6*, *Sox7* and *Sox17*, and mesodermal genes such as *Nodal*, *Edn1* and *Bmpr1a*, and down-regulated genes included pluripotency genes such as *Nanog, Pou5f1* and *Utf1* as well as cell cycle genes such as *Ccnd1*, *Ccne1*, *Esrrb* and *Rbbp4* (Gao et al., 2008; Kidder et al., 2011; Ostapcuk et al., 2018). In ESCs, BRG1, HP1γ and core pluripotency factors are known to bind the developmental genes to repress their expression, and occupy ESC-specific genes to support ESC pluripotency (Ho et al., 2009; Kidder et al., 2009; King and Klose, 2017; Mattout et al., 2015). In addition, esBAF members BRG1/BAF250/BAF155 and HP1γ are required for cell proliferation (Ho et al., 2008; Gao et al., 2008; Zaidan and Sridharan, 2020). Thus, similar to these factors, PBH is enriched at both highly-expressing ESC-specific genes that are down-regulated, and lowly-expressing PrE genes that are up-regulated during ESC differentiation, which strongly suggested that POGZ functions to maintain ESCs by association with BRG1 and HP1γ.

We have shown that POGZ, BRG1 and core pluripotency factors such as OCT4 are co-localized genome wide (Figure 4J-K). Next, we asked whether PBH co-occupy with NANOG and OCT4 genome wide. A Pearson correlation analysis of ChIP-seq signals revealed a high degree of co-localization among POGZ, BRG1, HP1, NANOG and OCT4 (Figure 6C). POGZ exhibited a considerable correlation with BRG1 (0.59). At a few ESC-specific and lineage specifying genes, OCT4/NANOG and PBH were co-localized at both proximal TSS and distal enhancers (Figure S6B). When extended this to genomic level, we found that PBH and OCT4/NANOG were extensively co-localized and shared a considerable number of target genes (about 1400) (Figure S6C). The results strongly suggested that PBH and OCT4/NANOG function together to maintain ESCs.

BRG1 is the core enzymatic components of the esBAF, which included many other subunits such as BAF155, BAF60a and SS18. The co-localization of BRG1 and POGZ prompted us to further investigate whether POGZ is co-localized with other members of the esBAF complex. To this end, we consulted the ChIP-seq data of SS18 and BAF155 (King and Klose, 2017; Ho et al., 2008). Global meta-analysis showed that POGZ, BRG1, BAF155 and SS18 were co-localized genome wide (Figure 6D). Snapshots of selected genes were shown for representatives (Figure 6E). Thus, we propose that POGZ co-localizes with and recruits esBAF genome wide in ESCs.

Finally, we examined the overlap of POGZ C&T peaks with repressive histone marks such as H3K9me3, H3K27me3 and H2Aub1. Global meta-analysis showed that POGZ was not significantly enriched with H3K9me3/H3K27me3/H2Aub1 (Figure 6F). Snapshots of selected genes clearly showed that POGZ exhibited strong signals at actively expressing *Pou5f1* gene decorated with H3K27ac/H3k4me1, and exhibited low signals at lowly expressing *Gata6* gene decorated with H2Aub1/H3K27me3 (Figure 6G). Taken together, we propose that POGZ was not significantly linked with heterochromatin regions decorated with H3K9me3, but with euchromatin in ESCs (Figure 4D; 6F).

### Nucleosome occupancy and positioning are altered in the absence of POGZ

The esBAF is a well known chromatin remodeler complex that regulates ESC chromatin state (Ho et al., 2008). Considering the close-relationship among POGZ, BRG1/esBAF, we asked whether loss of POGZ leads to alteration of chromatin accessibility, by performing ATAC-seq analysis for control and early passage *Pogz-/-* ESCs (passage number is 12).

A total of approximately 10,250 ATAC-seq peaks were identified in control ESCs. The majority of POGZ-bound loci were of ATAC-seq signals, suggesting that POGZ is bound to accessible chromatin or POGZ binding renders chromatin accessible (Figure S7A). Global meta-analysis showed that the chromatin accessibility was moderately reduced in the absence of POGZ, suggesting that POGZ primarily functions to promote chromatin accessibility (Figure 7A). Representative snapshots were shown for selected genes or chromosome fragment (Figure 7B; Figure S7A). In the absence of POGZ, chromatin accessibility was reduced at both TSS and enhancer regions (Figure 7B; Figure S4B; S7B). Bioinformatics analysis of POGZ-dependent ATAC-seq loci revealed that the top three motifs are the putative DNA motifs bound by pluripotency factors OCT4 and KLF4 (Figure S7C).

**Figure 7.**
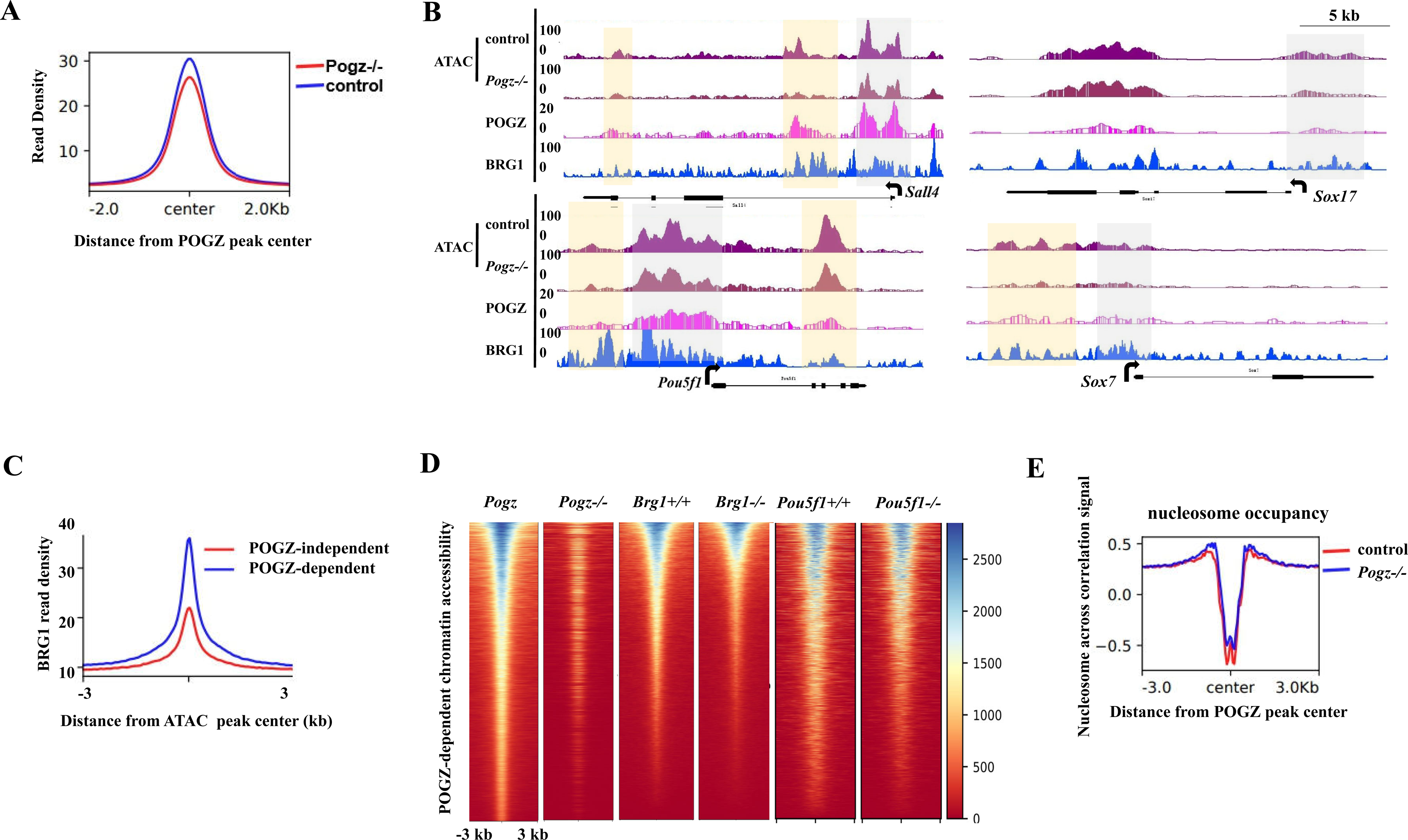
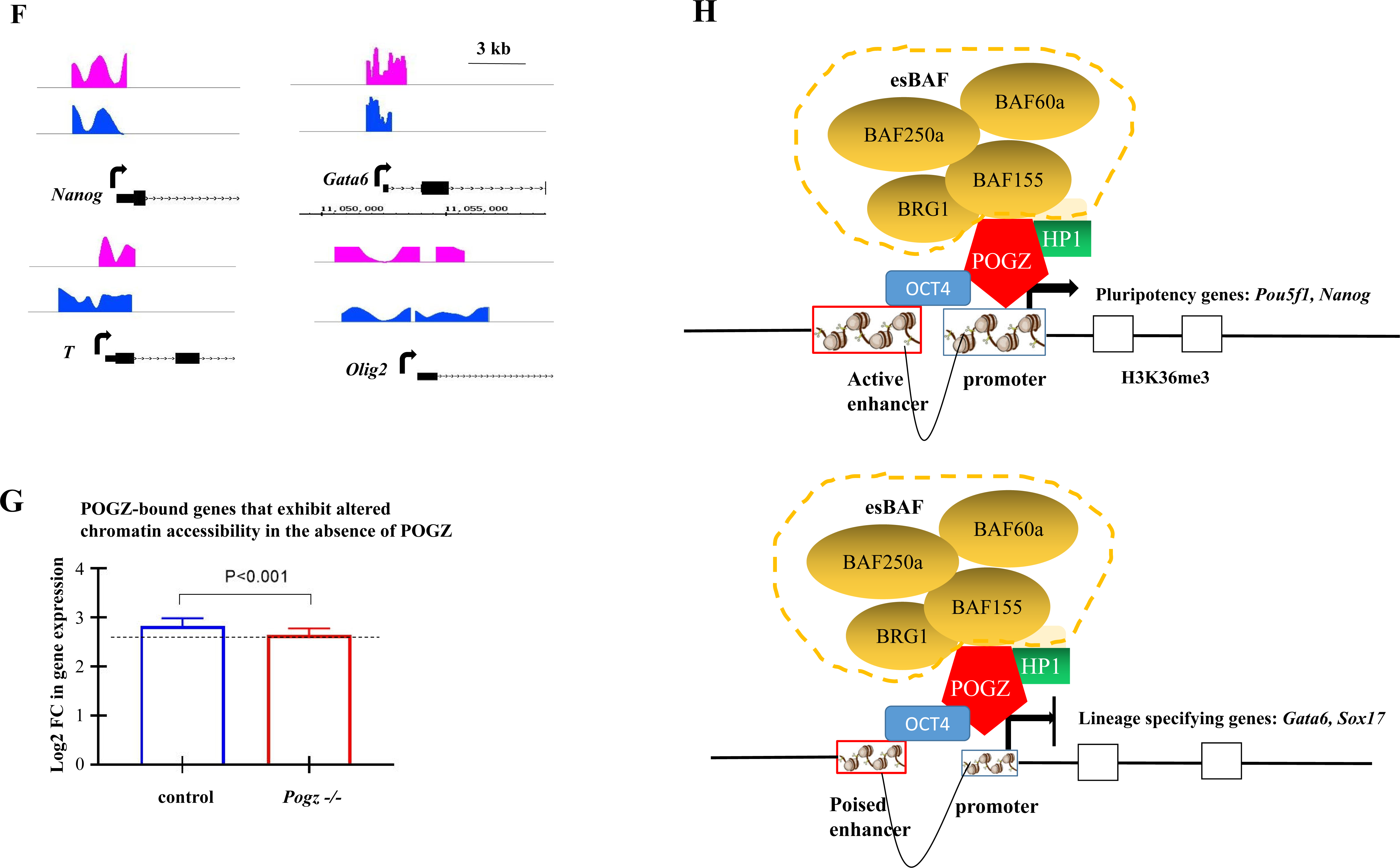
POGZ regulates chromatin accessibility by association with BRG1. **A** A metaplot showing the reduced chromatin accessibility at POGZ-peak center in *Pogz-/-* ESCs compared to control ESCs. **B** Snapshots of the indicated POGZ and BRG1 co-occupied loci showing that the chromatin accessibility was reduced in the absence of POGZ. Proximal TSS (grey) and distal enhancers (orange). **C** Metaplot showing that BRG1 was more enriched at sites where POGZ is responsible for chromatin accessibility. **D** Heat map of ATAC-seq signals at POGZ-dependent ATAC sites in control and BRG1-depleted ESCs, and in control and OCT4-depleted ESCs (King and Klose, 2017). Sites were ranked by loss of POGZ-dependent ATAC-seq signals (n=1630). Each row represents a 6 kb window centered on the peak midpoint. **E** Metaplot showing that nucleosome occupancy was slightly increased across sites bound by POGZ (n=16,728). **F** Close-up snapshots showing the change of nucleosome occupancy and phasing at the indicated gene loci. **G** Comparison analysis of expression levels for POGZ-bound genes that exhibited alteration in chromatin accessibility in control and *Pogz-/-* ESCs. **H** Cartoon showing the function of POGZ in ESCs. POGZ has a dual function: facilitating ESC-specific genes and suppressing PrE genes by recruiting BRG1/esBAF and HP1.

Next, we asked whether BRG1 is required for POGZ-regulated accessible chromatin by examining the overlap of BRG1 ChIP-seq peaks and POGZ-dependent ATAC peaks. We found that BRG1 is enriched at POGZ-dependent ATAC peaks (Figure 7B). Global meta-analysis showed that BRG1 ChIP signals were much stronger at sites where POGZ is responsible for chromatin accessibility (Figure 7C). This observation suggested that POGZ may require BRG1 to shape chromatin accessibility of POGZ-bound sites and gene regulation in the maintenance of ESC state. To investigate this, we consulted the published ATAC-seq data for BRG1 (King and Klose, 2017). When examining ATAC-seq signal at a few POGZ target sites in BRG1 knockout ESCs, we found a substantial reduction of chromatin accessibility, which was similar to that in *Pogz-/-* ESCs (Figure S7D). When extended to all POGZ-bound ATAC sites, a slight reduction of ATAC-seq signals was observed in BRG1-depleted ESCs, similar to *Pogz-/-* ESCs (Figure S7E). When examining POGZ-dependent ATAC sites, we found that ATAC-seq signals at these sites appeared more reduced in BRG1 knockout ESCs (Figure 7D). Thus, loss of BRG1 had a similar effect on chromatin accessibility to that by loss of POGZ.

As PBH co-occupy with OCT4 genome wide, and OCT4 requires BRG1 to establish an ESC-specific chromatin accessibility (King and Klose, 2017), we speculated that loss of either OCT4 or POGZ may lead to a similar change of chromatin accessibility. To investigate this, we downloaded the published ATAC-seq data for OCT4 (King and Klose, 2017). When examining ATAC-seq signal at a few POGZ target sites in OCT4 knockout ESCs, we found a substantial reduction of chromatin accessibility (Figure S7D). When extended to all POGZ-dependent ATAC sites, a slight reduction of ATAC-seq signals was observed in OCT4-depleted ESCs, similar to BRG1-depleted ESCs (Figure 7D).

Next, we asked whether POGZ regulates nucleosome occupancy at target genes. Global meta-analysis showed that loss of POGZ at POGZ-bound sites led to an increase of nucleosome occupancy (Figure 7E). When a few ESC-specific and lineage-specifying genes were examined, alteration of both nucleosome occupancy and phasing was observed (Figure 7F).

Finally, we investigated whether the alteration of chromatin accessibility is relevant to gene regulation by loss of POGZ. At a global level, POGZ-bound genes which exhibited altered ATAC-seq signals in the absence of POGZ was significantly down-regulated (Figure 7G), thus establishing a correlation between gene expression and chromatin accessibility. Nevertheless, we found that many genes, which exhibited dramatic change in chromatin accessibility in the absence of POGZ, were not deregulated at all. It appeared that alteration of chromatin accessibility of POGZ-bound genes did not always lead to gene expression change. This was also observed in OCT4- and ADNP-depleted ESCs where much more dramatic change in chromatin accessibility occurred (King and Klose, 2017; Ostapcuk et al., 2018; Sun et al., 2020).

## Discussion

Our findings were surprising at first glance as POGZ has been shown to interact with HP1s, which reads the repressive H3K9me3 histone mark that is known to be associated with transcriptional repression (Vermeulen et al., 2010). POGZ was also shown to function as a negative regulator of transcription based on an in vitro GAL4-DBD luciferase assay (Suliman-Lavie et al., 2020). In addition, POGZ is found frequently linked with factors such as ADNP, MGA and CHD4 which are known transcriptional repressors (Ostapcuk et al., 2018). In this work, using biochemistry, bioinformatics and functional analyses, we found that POGZ is closely linked with esBAF complex, and is localized to promoter and enhancer regions. In fact, we found that POGZ is enriched to H3K4me1 and H3K27ac marked active enhancers and super-enhancers in ESCs (Markenscoff-Papadimitriou et al., 2021). POGZ can function as both a transcriptional activator and repressor, similar to the core pluripotency factors and BRG1/esBAF: facilitating ESC-specific genes and suppressing lineage-specifying genes (Figure 7H). Consistently, loss of POGZ leads to significant down-regulation of pluripotency-associated genes, and up-regulation of differentiation genes, which eventually drives ESCs exit from pluripotency.

POGZ has been shown to be essential for normal mitotic progression of HeLa cells (Nozawa et al., 2010). We found that *Pogz-/-* ESCs are difficult to passage and eventually collapse around passage 20, accompanied by gradual loss of stemness, reduced cell proliferation and apoptosis. In fact, depletion of BRG1 and HP1γ has been shown to lead to similar phenotypes in ESCs. Consistently, PBH binds pluripotency and cell cycle genes which are down-regulated in the absence of each factor (Figure 6B). In this work, we performed all the differentiation, ChIP-seq and ATAC-seq experiments using early passage *Pogz-/-* ESCs (mostly at passage 12-14) where OCT4 and/or NANOG are abundantly expressed (Figure 1D).

*POGZ* is one of the top recurrently mutated genes in patients with NDDs, particularly ASD and ID. In addition to neurodevelopmental defects, *POGZ* patients may exhibit additional deficits, such as short stature, cardiac problem, hypotonia, strabismus, hearing loss, and abnormal craniofacial formation, including brachycephaly, long and flat malar region, broad nasal tip, short philtrum and thin vermillion border (White et al., 2016). The molecular and cellular mechanisms underlying the pleiotropic phenotypes by *POGZ* mutation remain unclear. Our work using ESC model leads to several important observations. First, POGZ might be a master regulator of neural development. Both GO and KEGG analyses indicated that POGZ target genes are overwhelmingly linked with neuronal function and neurodevelopmental disorders. This was in line with that POGZ is one of the most recurrently mutated genes in patients with neurodevelopmental disorders. Second, POGZ associates and recruits esBAF and HP1s genome wide. As esBAF complex and HP1s can control the expression of target genes directly and indirectly, by modulating the chromatin and epigenome, it is not unexpected that various genes including pluripotency-, endodermal- and mesodermal genes as well as signaling components were deregulated in the absence of POGZ. Third, POGZ is an important pluripotency-associated factor. There is an extensive overlap of PBH and NANOG/OCT4 peaks genome-wide, and POGZ interacts with OCT4. POGZ binding motifs are similar to those bound by OCT4 and KLF4. PBH is required to promote ESC-specific genes, and repress PrE genes, similar to OCT4/NANOG. Consistently, loss of POGZ leads to a similar chromatin change to loss of OCT4, and POGZ-associated BRG1 and HP1s are essential regulators of ESCs (King and Klose, 2017; Mattout et al., 2015; Zaidan and Sridharan, 2020). Fourth, many imprinted genes were deregulated in the absence of POGZ. Imprinted genes are known to be linked to a number of human behavioral and neural developmental disorders (leung et al., 2011). This suggested that loss of POGZ may also contribute to NDDs by increasing the expression of imprinted genes. A few imprinted genes such as *Nespas* and *Rian* have been shown to be deregulated in cortex of *Pogz-/-* mice (Markenscoff-Papadimitriou et al., 2021). Taken together, it can be concluded that POGZ is a multifunctional protein that is involved in the regulation of signaling pathways, chromatin structuring, genome imprinting and epigenome. Dysfunction in any of these processes can lead to gene expression and developmental defects, which may explain the pleiotropic phenotypes observed in *POGZ* patients.

How the different chromatin remodeler complexes and HP1s are localized to specific genome loci is an important yet unsolved question (Ho et al., 2009; Zaidan and Sridharan, 2020). ADNP has been shown to recruit CHD4 and HP1s to the target sites in a DNA sequence specific manner (Ostapcuk et al., 2018). In this work, we show that POGZ recruits BRG1 and HP1s to POGZ-bound sites in a similar way. Although lack of a global ChIP-seq analysis and quantitative assay of PBH triplex as described by Ostapcuk (2018), our biochemistry and functional experiments strongly suggested that POGZ associates with and recruits BRG1 (therefore esBAF) and HP1s to the target sites. Thus, work by us and others suggest that DNA-binding TFs, particularly the C2H2-type zinc finger of transcription factors, are likely important players in recruiting different chromatin remodeler complexes to specific genome loci. It will be interesting to investigate this further in future.

## Materials and Methods

### ES Cell Culture

Mouse embryonic stem cells (mESCs) R1 were maintained in Dulbecco’s Modified Eagle Medium (DMEM, BI, 01-052-1ACS) containing 20% knockout serum replacement (KSR, Gibco, 10828028), 1 mM sodium pyruvate (Sigma, S8636), 2 mM L-Glutamine (Sigma, G7513), 1,000 U/ml leukemia inhibitory factor (LIF, Millipore, ESG1107,) and penicillin/streptomycin (Gibco, 15140-122) at 37°C with 5% CO2. Cells were routinely propagated by trypsinization and replated every 3-4 days, with a split ratio of 1: 6.

### Embryoid body formation

ESCs differentiation into embryoid bodies (EBs) was performed in attachment or suspension culture in medium lacking LIF and knockout serum replacement (KSR), as described previously (Sun et al., 2020).

### Generation of *Pogz-/-* ESCs

*Pogz-/-* mESCs were generated by CRISPR/Cas9 technology (Ran et al., 2013). Briefly, we designed gRNAs on exon 2 of *Pogz* gene by using the online website http://crispr.mit.edu/. The gRNA sequences are 5’-CGACCTGTTTATGGAATGTGAGG-3’. The guide sequence oligos were cloned into a plasmid containing Cas9 and the sgRNA scaffold (pSpCas9(BB)) with a puromycin resistance gene (Ran et al., 2013). After 48 hours of targeting plasmids transfection, mESCs were selected with 1-2 μg/ml puromycin for 7 days. Then the cells were re-seeded on a 10-cm dish coated with 0.1% gelatin to form colonies. The single colony was picked up, trypsinized and passaged at low density. DNA from single colonies from the passaged cells was extracted and used for genotyping.

### Generation of 3×Flag-POGZ restoring *Pogz-/-* mESC Cell Lines

The full-length *Pogz* cDNA (NM_172683.4) was amplified by PCR and then cloned into pCMV-3×Flag vector. The full-length *Pogz* cDNA sequence containing N-terminal 3×Flag sequence was subcloned into pCAG-IRES-Puro vector. To make stable transgenic cells, *Pogz-/-* mESCs were transfected with pCAG-IRES-Puro-3×Flag-*Pogz* vector using Lipofectamine 2000 (Gibco, 11668019).

48 hours later, cells were selected by 1 μg/ml puromycin. After 4-5 days drug selection, cells were expanded and passaged. Western Blot assay was performed to confirm the transgenic cell line using FLAG antibodies.

### RNA preparation, RT-qPCR and RNA-seq

Total RNA from mESCs and ESC-derived EBs was extracted with a Total RNA kit (Omega, R6834-01). A total of 1 μg RNA was reverse transcribed into cDNA using the TransScript All-in-One First-Strand cDNA synthesis Supermix (Transgen Biotech, China, AT341). Quantitative real-time PCR (qRT-PCR) was performed using the TransStart® Tip Green qPCR SuperMix (Transgen Biotech, China, AQ-141). The primers used for qRT-PCR were previously described (Sun et al. 2020). All experiments were repeated for at least three times. The relative gene expression levels were calculated based on the 2^-ΔΔCt^ method. Data were shown as means ± S.D. The Student’s t test was used for the statistical analysis. The significance is indicated as follows: *, *p* < 0.05; **, *p* < 0.01; ***, *p* < 0.001.

For RNA-seq, control and mutant ESCs, and day 6 control and *Pogz-/-* ESC-derived EBs were collected and treated with Trizol (Invitrogen). RNAs were quantified by a Nanodrop instrument, and sent to BGI Shenzhen (Wuhan, China) for making RNA-seq libraries and deep sequencing. For each time points, at least two biological repeats were sequenced. DEGs were defined by FDR*<* 0.05 and a Log2 fold change*>* 1 was deemed to be differentially expressed genes (DEGs). Less strictly, genes whose expression differed by > 1.5 with a p-value < 0.05 were deemed differentially expresssed genes (DEGs).

We consulted RNA-seq for HP1 KO ESCs (GSM2582351-2582353; GSM2582375), for *Brg1-/-* ESCs (GSM2341328-2341333; GSM734277-79) (King and Klose, 2017; Ostapcuk et al., 2018; Sridharan et al., 2013).

### Protein extraction and Western blot analysis

For protein extraction, ESCs or HEK293T cells were harvested and lysed in TEN buffer (50 mM Tris-HCl, 150 mM NaCl, 5 mM EDTA, 1% Triton X-100, 0.5% Na-Deoxycholate, supplement with Roche cOmplete Protease Inhibitor). The lysates were quantified by the Bradford method and equal amount of proteins were loaded for Western blot assay. Antibodies used for WB were POGZ (Abcam, ab171934), anti-BRG1 (Proteintech, 21634-1-AP), anti-HP1gamma (Proteintech, 11650-2-AP), anti-CHD4 (Proteintech, 14173-1-AP), anti-OCT4 (Proteintech, 60242-1-Ig), Anti-FLAG (F1804/F3165, Sigma, 1: 1000), anti-MYC antibody (Transgen Biotech, HT101) and anti-HA (Abbkine, A02040, 1: 1000). Briefly, the proteins were separated by 10% SDS-PAGE and transferred to a PVDF membrane. After blocking with 5% (w/v) non-fat milk for 1 hour at room temperature, the membrane was incubated overnight at 4°C with the primary antibodies. Then the membranes were incubated with a HRP-conjugated goat anti-rabbit IgG (GtxRb-003-DHRPX, ImmunoReagents, 1: 5000), a HRP-linked anti-mouse IgG (7076S, Cell Signaling Technology, 1: 5000) for 1 hour at room temperature. The GE ImageQuant LAS4000 mini luminescent image analyzer was used for photographing. Western blot experiments were repeated at least two times.

Quantification of Western blot bands was performed by ImageJ software, according to the website: https://imagej.nih.gov/ij/docs/guide/146-30.html. Briefly, the rectangle tool was selected and used to draw a box around the lane, making sure to include some of the empty gel between lanes and white space outside of the band. All lanes were selected one by one. Once all lanes are defined, go to Plot lanes to generate histograms of each lane. Then the relative values were calculated by dividing each value by the control lane. The value of the control bands was set at 1.

### Co-immunoprecipitation assay (co-IP)

Co-IPs were performed with the Dynabeads Protein G (Life Technologies, 10004D) according to the manufacturer’s instructions. Briefly, 1.5 mg Dynabeads was conjugated with antibodies or IgG overnight at 4°C. Antibodies were used are: 10 μg IgG (Proteintech, B900610), or 10 μg anti-POGZ antibody, or 10 μg anti-FLAG antibody (Sigma, F3165/F1804), or 10 μg anti-HA antibody (Abbkine, A02040) or 10 μg anti-MYC antibody (Transgen Biotech, HT101). The next day, total cell lysates and the antibody-conjugated Dynabeads were incubated overnight at 4°C with shaking. After three-times washing with PBS containing 0.1% Tween, the beads were boiled at 95°C for 5 minutes with the 6×Protein loading buffer and the supernatant was collected for future WB analysis.

### Immunofluorescence assay

ESCs or ESC-derived cells were collected and fixed with 4% paraformaldehyde for half an hour at room temperature. Then the cells were washed with PBST (phosphate-buffered saline, 0.1% Triton X-100) for three times, each for 15 minutes. Following the incubation with blocking buffer (5% normal horse serum, 0.1% Triton X-100, in PBS) for 2 hours at room temperature, the cells were incubated with primary antibodies at 4°C overnight. Antibodies used were OCT4 (Proteintech, 60242-1-Ig), NANOG (Bethyl, A300-397A), GATA6 (Abcam, ab175349), HP1 (Abcam, ab213167), PH3 (CST, #9701). After three-times wash with PBST, the cells were incubated with secondary antibodies (1: 500 dilution in blocking buffer, Alexa Fluor 488, Life Technologies) at room temperature for 1 hour in the dark. The nuclei were counter-stained with DAPI (Sigma, D9542, 1: 1000). After washing with PBS for twice, the slides were mounted with 100% glycerol on histological slides. Images were taken by a Leica SP8 laser scanning confocal microscope (Wetzlar, Germany). About 10 images were taken for each slide. Quantification of IF staining was performed as described (Sun et al., 2020).

### Immunoprecipitation in combination with mass spectrometry

For Mass Spectrometry, the IP samples (immunoprecipitated by IgG or POGZ antibody) were run on SDS-PAGE gels and stained with the Coomassie Blue. Then the entire lanes for each IP samples were cut off and transferred into a 15 ml tube containing deionized water. The treatment of the samples and the Mass Spectrometry analysis were done by GeneCreate Biological Engineering Company (Wuhan, China).

### Protein-protein interaction assay using a rabbit reticulocyte lysate system

Protein-protein interaction assay using a rabbit reticulocyte lysate system has been described previously (Sun et al., 2020). Tagged-POGZ, Tagged-POGZ mutants, Tagged-BRG1 and Tagged-BRG1 mutants, as well as Tagged-BAF15 were synthesized using the TNT coupled reticulocyte lysate system according to the manual (Promega, L5020, USA). Briefly, 1 μg of circular PCS2- version of plasmids were added directly to the TNT lysates and incubated for 1.5 hours at 30°C. 1 μl of the reaction products were subjected to WB assay to evaluate the synthesized protein.

For protein-protein interaction assay, 5-10 μl of the synthesized HA or FLAG tagged proteins were mixed in a 1.5 ml tube loaded with the 300 μl TEN buffer, and the mixture was shaken for 30 minutes at room temperature. Next, IP or pull-down assay was performed using Dynabeads protein G coupled with anti-FLAG or anti-HA antibodies as described above.

### Apoptosis Analysis by Flow Cytometry

Annexin V-EGFP/PI (Propidium Iodide) Apoptosis Detection Kit (Yeason, 40303ES20) was used to detect apoptotic cells according to the manual. Briefly, ESCs were treated with trypsin without EDTA, and then washed twice by DPBS. Then cells were washed with 1 × Binding Buffer and incubated with Annexin V-EGFP for 5 minutes at room temperature in the dark. Then cells were treated with PI Staining Solution. Finally, apoptotic cells were detected by flow cytometry (BD, AccuriC6). The late apoptosis and necrotic cells will be AnnexinV^+^/PI^+^. The results were analyzed by Wan Yan from the Analysis and Testing Center of Institute of Hydrobiology, Wuhan.

### Quantification and statistical analysis

Data are presented as mean values ±SD unless otherwise stated. Data were analyzed using Student’s t-test analysis. Error bars represent s.e.m. Differences in means were statistically significant when *p*< 0.05. Significant levels are: \**p*< 0.05; \*\**P*< 0.01, and \*\*\**P*< 0.001.

## Bioinformatics

### ChIP-seq analysis

ChIP-seq data were aligned in Bowtie2 (version 2.2.5) with default settings. Non-aligning and multiple mappers were filtered out. Peaks were called on replicates using the corresponding inputs as background. MACS2 (version 2.1.1) was run with the default parameters. Peaks detected in at least two out of three replicates were kept for further analysis. BigWig files displaying the full length for uniquely mapping reads were generated using the bedGraphToBigWig (UCSC binary utilities).

To investigate the co-occupancy of POGZ, BRG1 and HP1γ/CBX3, we consulted previously published ChIP-seq data sets for BRG1 (GSE87820) and HP1γ/CBX3 (GSM1081158) (Ostapcuk et al., 2018; King and Klose, 2017; Sridharan et al., 2013). To investigate the co-occupancy of POGZ with H3K4me1 and H3K27ac, we consulted previously published ChIP-seq data sets for H3K4me1 (GSM2575694) and H3K27Ac (GSM2575695) (Dieuleveult et al., 2016). KLF4 (GSM4072779), ESRRB (GSM4087822), OCT4 (GSM2341284), NANOG (GSM2521520), MED1 (GSM4060038), SS18 (GSM2521508), BAF250a (GSM3318684), BAF155 (GSE69140), H3K27me3 (GSM4303791), H2Aub1 (GSM2865672) ChIP-seq data were also downloaded for this work.

### ATAC-seq Analysis

Paired-end reads were aligned using Bowtie2 using default parameters. Only uniquely mapping reads were kept for further analysis. These uniquely mapping reads were used to generate bigwig genome coverage files similar to ChIP–seq. Heat maps were generated using deeptools2. For the meta-profiles, the average fragment count per 10-bp bin was normalized to the mean fragment count in the first and last five bins, which ensures that the background signal is set to one for all experiments. Merged ATAC-seq datasets were used to extract signal corresponding to nucleosome occupancy information with NucleoATAC.

For comparison analysis of POGZ, OCT4 and BRG1 ATAC-seq signals, we downloaded previously published ATAC-seq data sets from Brg1 KD ESCs (GSM1941485-6), Brg1 KO ESCs (GSM2341280) and OCT4 KO ESCs (GSE87819) (Dieuleveult et al., 2016; King and Klose, 2017).

### Differential Binding And Gene Expression Analysis

Significant changes in ATAC-seq were identified using the DiffBind package, a FDR < 0.05 and log2 fold change> 1 was deemed to be a significant change. Gene ontology (GO) analysis for differentially regulated genes, and heat maps were generated from averaged replicates using the command line version of deepTools2. Peak centers were calculated based on the peak regions identified by MACS (see above).

### Data availability

All RNA-seq, ATAC-seq, ChIP-seq and CUT&Tag data have been deposited in the public database at Beijing Genomic Institute (BGI) at https://bigd.big.ac.cn/, with the accession number of CRA003852.

## Acknowledgments

This work was supported by National Key Research and Development Program (2016YFA0101100), and National Natural Science Foundation of China (31671526) to YH Sun.

## Author contributions

XY Sun performed the experiments; LX Cheng performed the bioinformatics analysis; YH Sun designed the work, provided the final support of the work, and wrote the paper.

---

**Figure S1.**
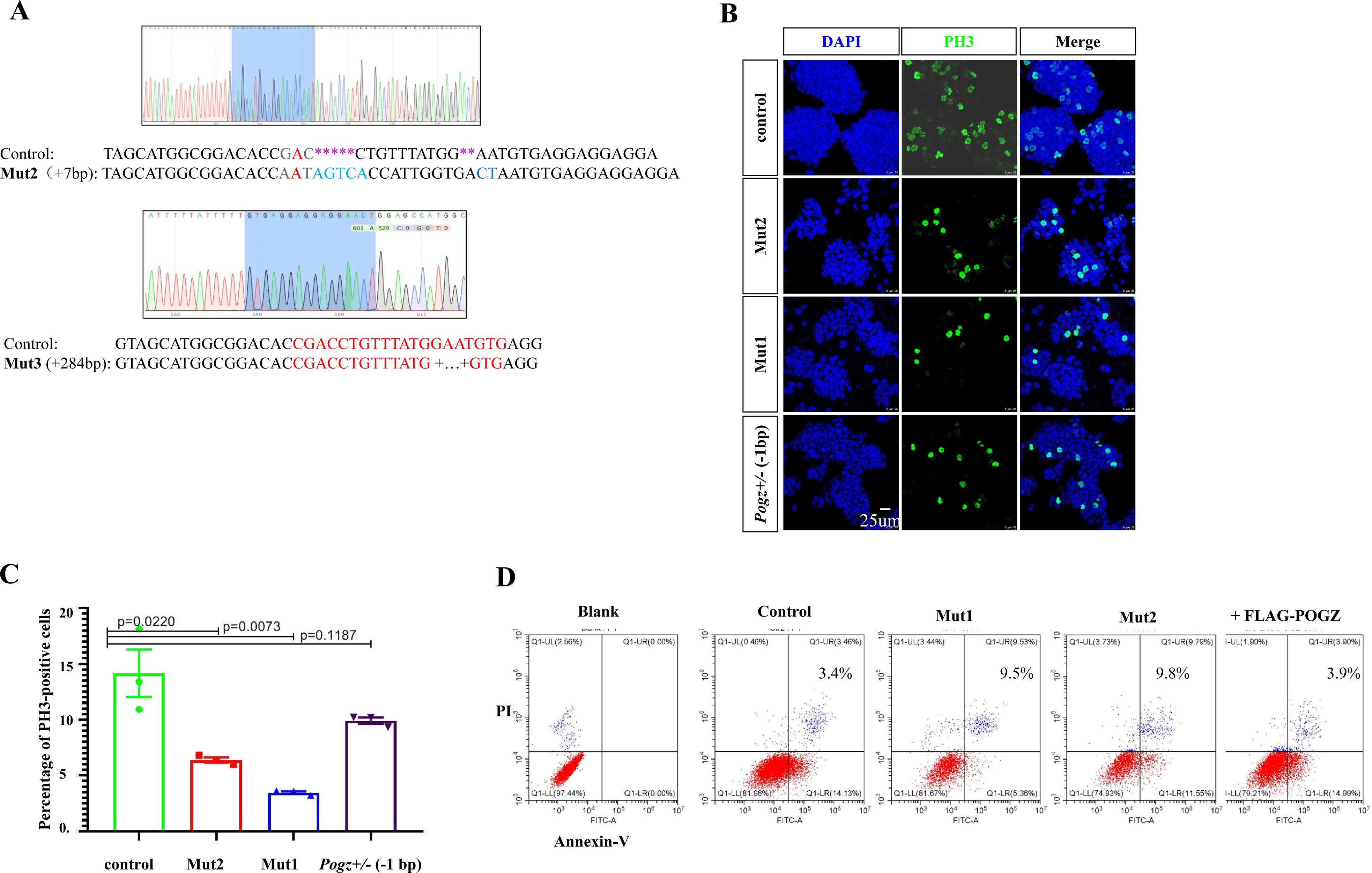
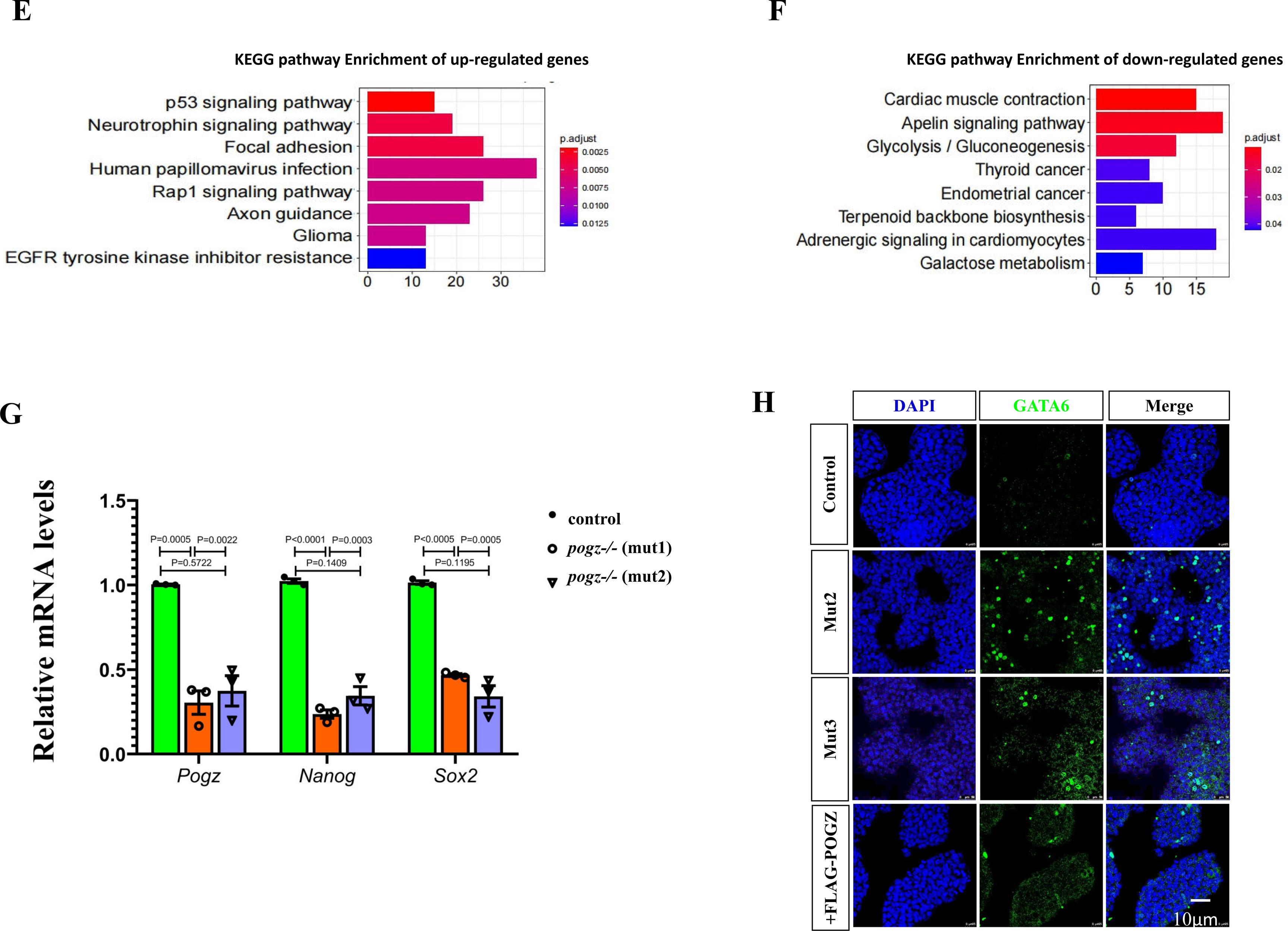
Related to Figure 1. **A** Genotyping showing two additional mutant alleles: Mut2 and 3. **B** PH3 staining of control ESCs, Mut1- and Mut2-*Pogz^-/-^* and *Pogz^+/-^* ESCs at passage 12, showing reduced number of PH3 positive cells in homologous and heterozygous mutant ESCs. **C** Quantification of (B). **D** Apoptosis analysis by flow cytometry for control-, Mut1- (passage 12), Mut2- (passage 12), and FLAG-POGZ restoring ESCs. **E** KEGG analysis of up-regulated DEGs between control and *Pogz*^-/-^ ESCs. **H** KEGG analysis of down-regulated DEGs between control and *Pogz^-/-^* ESCs. **G** qRT-PCR results for the indicated genes of control, Mut1 and Mut2 ESCs at passage 18. **H** IF staining results showing that GATA6 was abnormally expressed in Mut2 and Mut3 *Pogz*^-/-^ ESCs (passage number is 18). Bar: 25 μm.

**Supplementary Figure 2.**
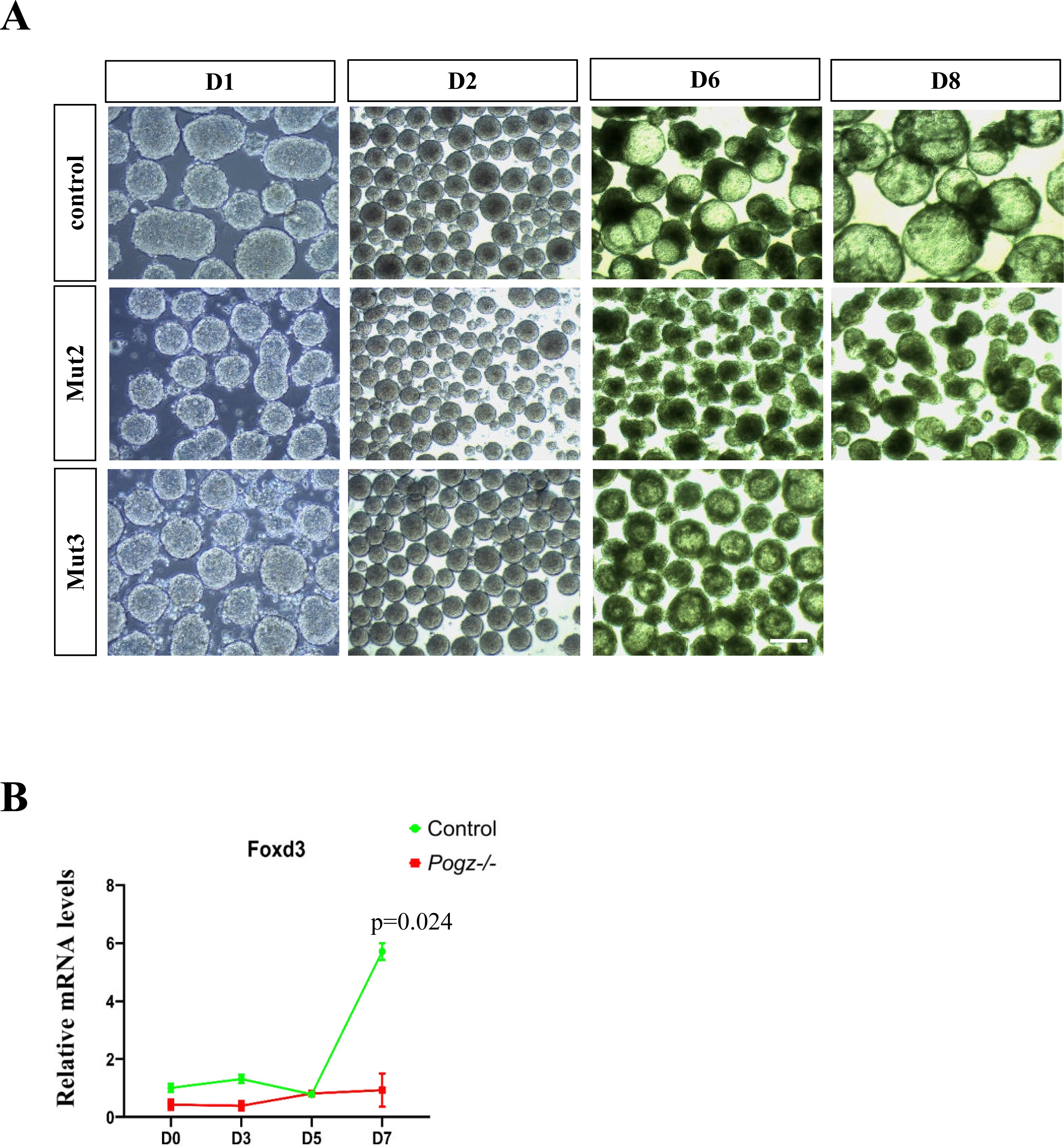
Related to Figure 2. **A** Morphology of day 1, 2, 6 and 8 EBs from control and two additional *Pogz* mutant ESCs (Mut2 and 3 at passage 12). **B** Time course qRT-PCR analysis of *Foxd3* during control and Mut1 ESC directional differentiation towards neural progenitors. Bar: 25 μm.

**Figure S3.**
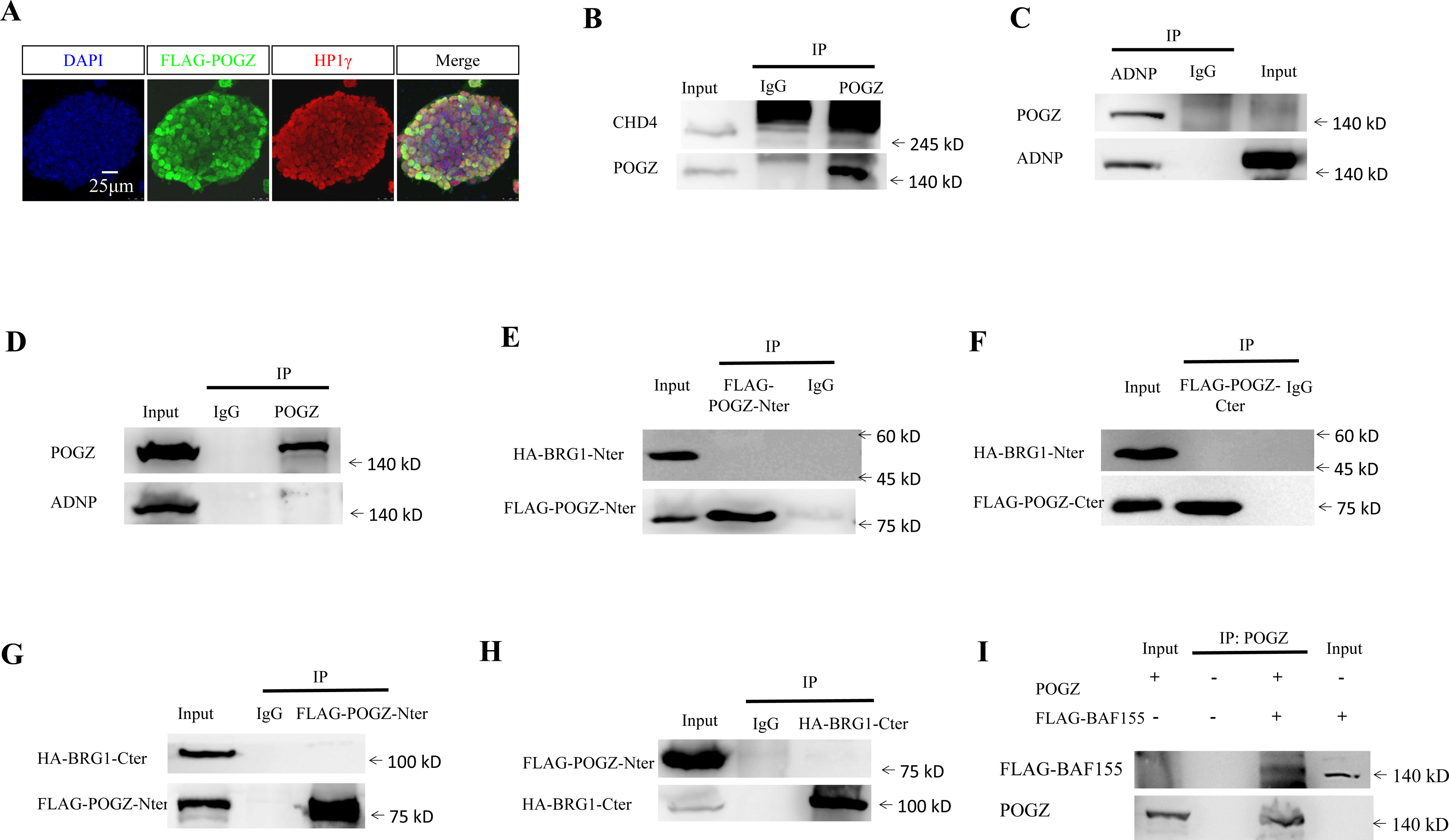
Related to Figure 3. **A** Double IF staining of HP1γ and FLAG-POGZ. **B** IP results showing that POGZ failed to interact with CHD4 in ESCs. **C** Co-IP results showing that ADNP failed to interact with POGZ in ESCs. **D** Co-IP results showing that POGZ failed to interact with ADNP in ESCs. **E** IP results showing FLAG-POGZ-Nter failed to pull down HA-BRG1-Nter in 293T cells. **F** IP results showing FLAG-POGZ-Cter failed to pull down HA-BRG1-Nter in 293T cells. **G** IP results showing FLAG-POGZ-Nter failed to pull down HA-BRG1-Cter in 293T cells. **H** IP results showing HA-BRG1-Cter failed to pull down FLAG-POGZ-Nter in 293T cells. **I** Direct interaction of in vitro synthesized POGZ and FLAG-BAF155 by TnT system. IP: POGZ, WB: FLAG. All experiments were repeated at least two times, and shown are representative images. Bar: 10 μm.

**Figure S4.**
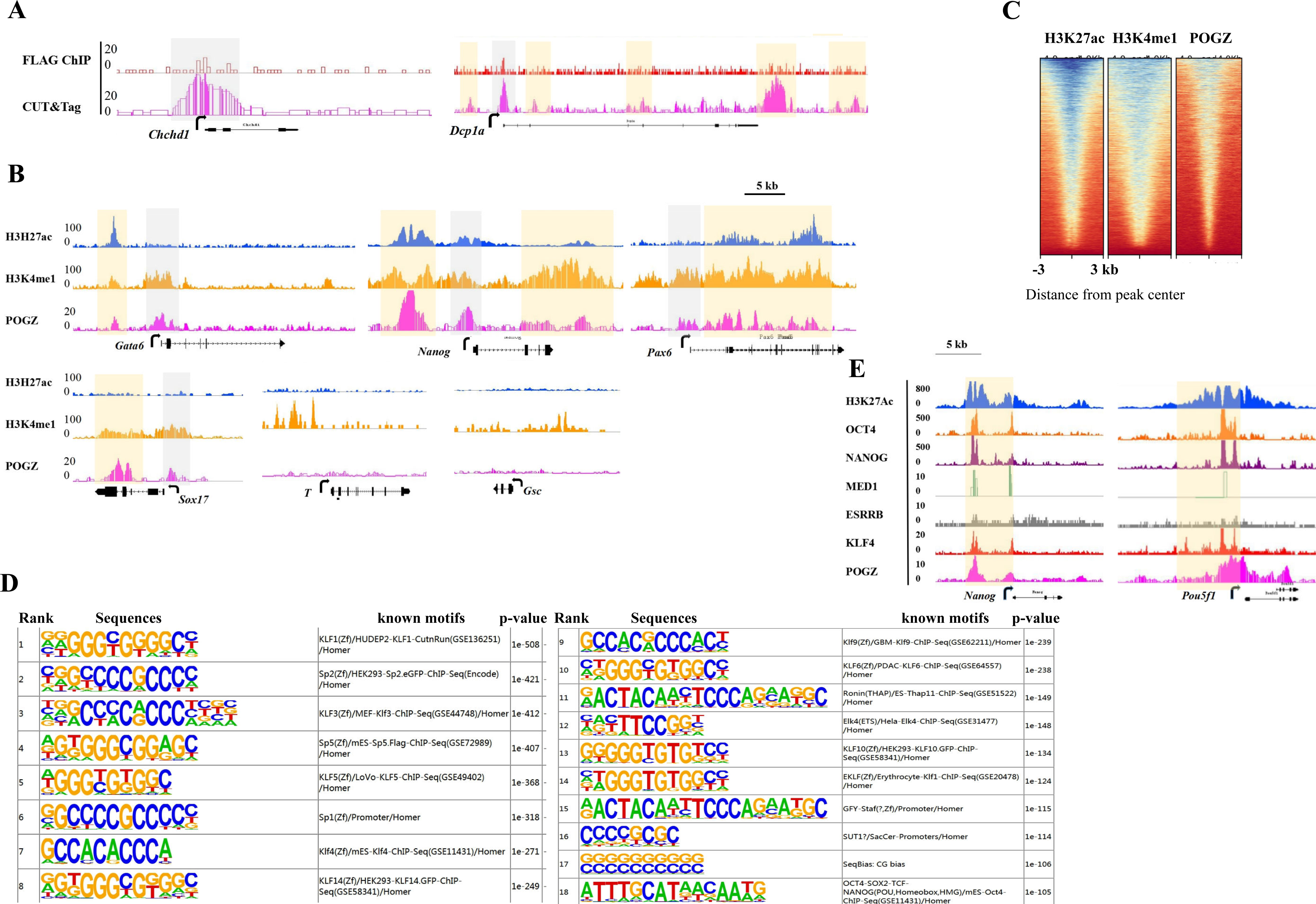
Related to Figure 4. **A** Snapshots of ChIP and CUT&Tag signals at *Chchd1* and *Dcp1a* loci, showing that CUT&Tag outperforms FLAG ChIP-seq. Grey: proximal TSS; Orange: gene distal or gene body regions. **B** Snapshots ChIP and CUT&Tag signals at the indicated loci, showing that POGZ is localized to gene loci decorated with H3K4me1 and H3K27Ac, respectively. Grey: proximal TSS; Orange: enhancer regions. Note the broad POGZ signals at *Nanog*, *Sox17* and *Pax6* loci, and low signals at *T* and *Gsc*. **C** Heat map of H3K27ac, H3K4me1 and POGZ enrichment across sites (n=8500) bound by both H3K27ac and H3K4me1 (therefore mark active enhancers). Each row represents a 6 kb window centered on the peak midpoint. **D** A rank of motifs identified in POGZ binding sites by HOMER. Shown were the top 18 motifs with p-value. **E** Snapshots of ChIP and CUT&Tag signals at *Nanog* and *Pou5f1* loci, showing a broad overlapping occupancy of POGZ, OCT4, NANOG, H3K27Ac, ESRRB, KLF4 and MED1, indicating that POGZ is localized to super-enhancers. Orange: enhancer regions.

**Figure S5.**
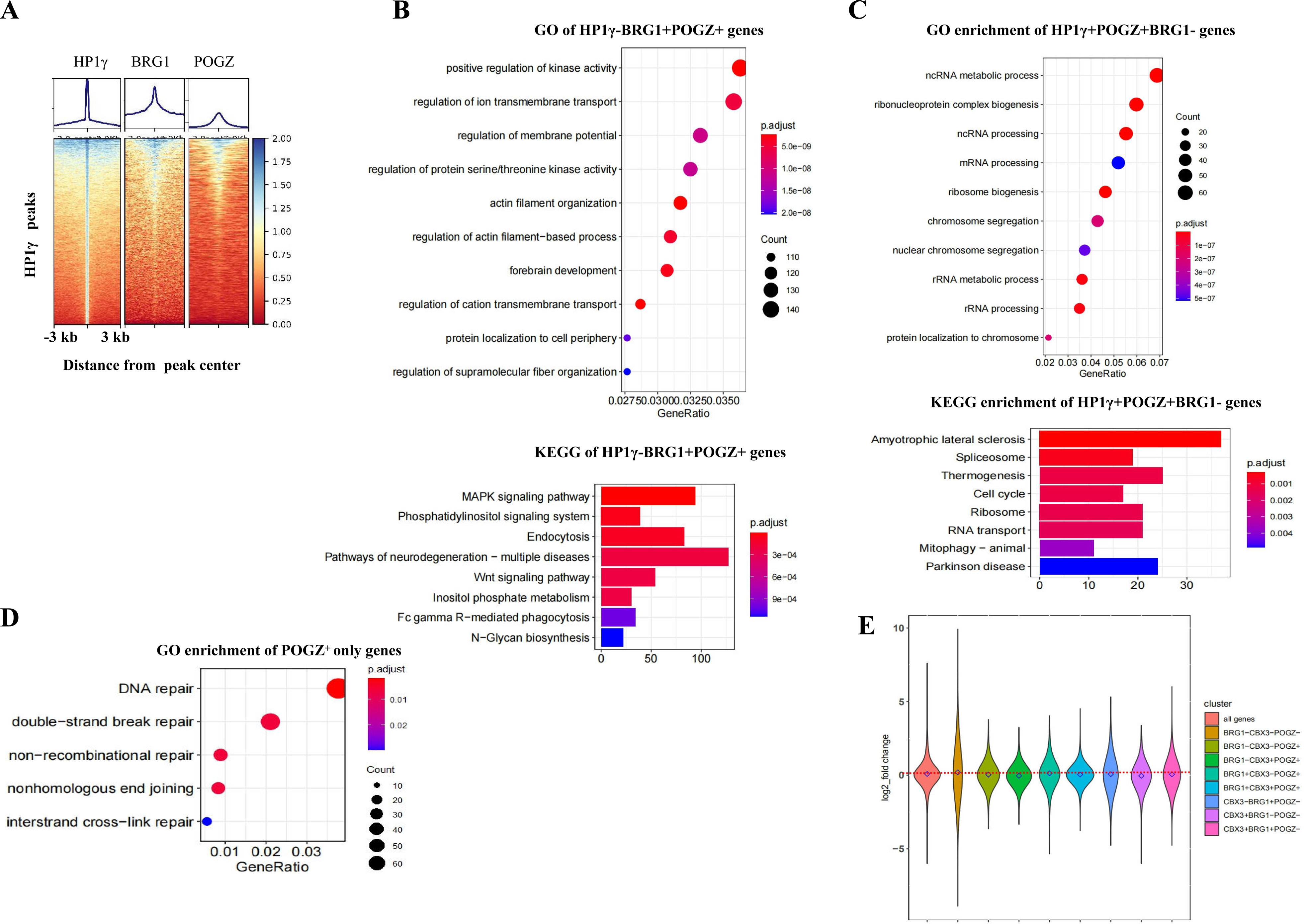
Related to Figure 5. **A** Heat map of BRG1, HP1γ and POGZ enrichment across sites bound by HP1γ (n=2156). Each row represents a 6 kb window centered on the peak midpoint. **B** (Up) GO analysis of HP1γ-BRG1+POGZ+ genes; (Bottom) KEGG analysis of HP1γ-BRG1+POGZ+ genes. **C** (Up) GO analysis of HP1γ+BRG1-POGZ+ genes; (Bottom) KEGG analysis of HP1γ+BRG1-POGZ+ genes. **D** GO analysis of HP1γ-BRG1-POGZ+ genes. **E** Change of average expression levels of each gene cluster, compared to that of all genes, in the absence of POGZ.

**Figure S6.**
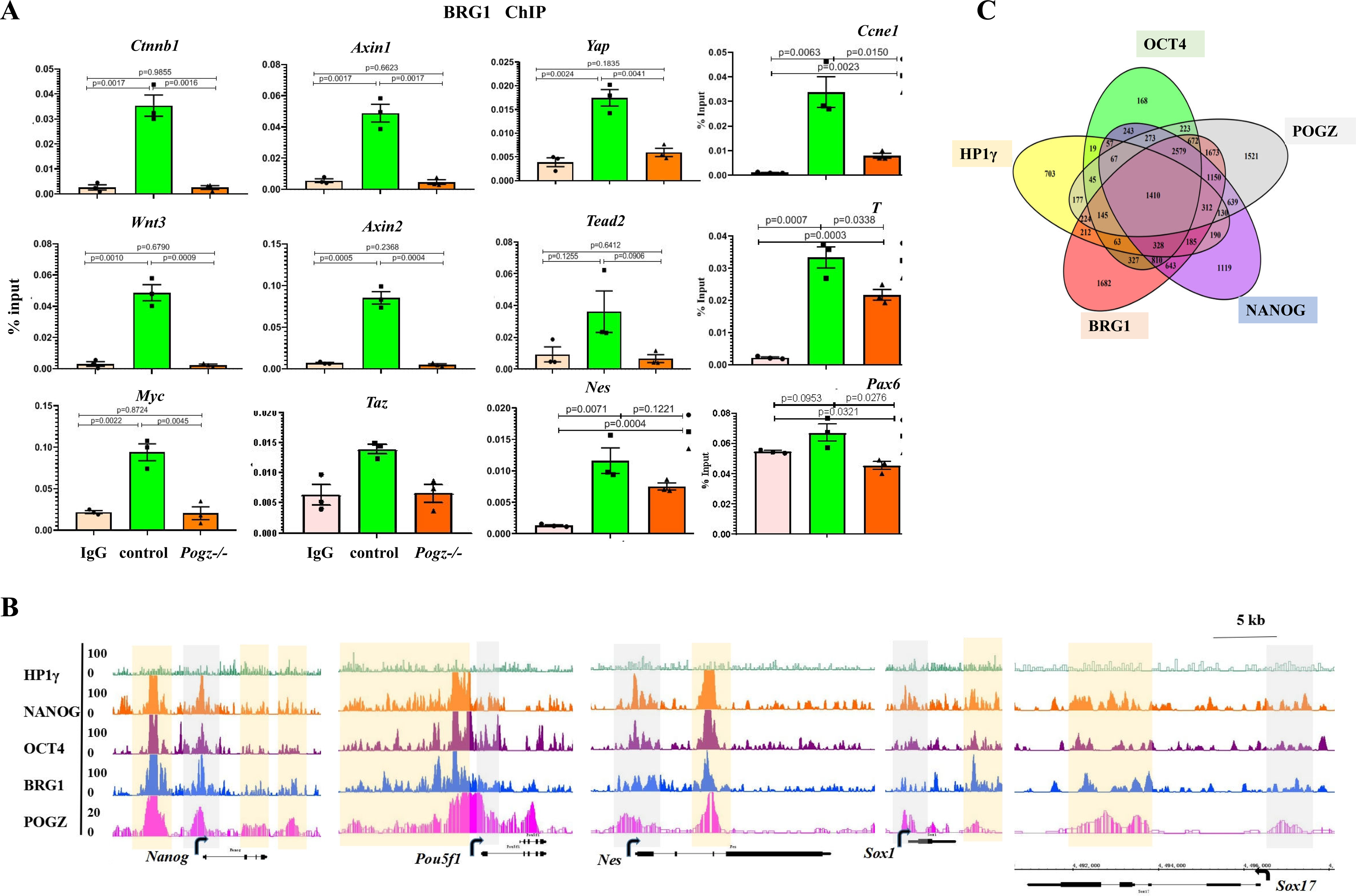
**A** ChIP-PCR results showing that BRG1 levels were reduced at the indicated genes, in the absence of POGZ. **B** ChIP-seq snapshots showing the co-localization of POGZ, BRG1, HP1γ/CBX3, NANOG and OCT4 at the indicated gene loci. TSS (grey color) and distal enhancer regions (orange). **C** Venn diagram showing the shared targets among POGZ, BRG1, HP1γ/CBX3, NANOG and OCT4.

**Figure S7.**
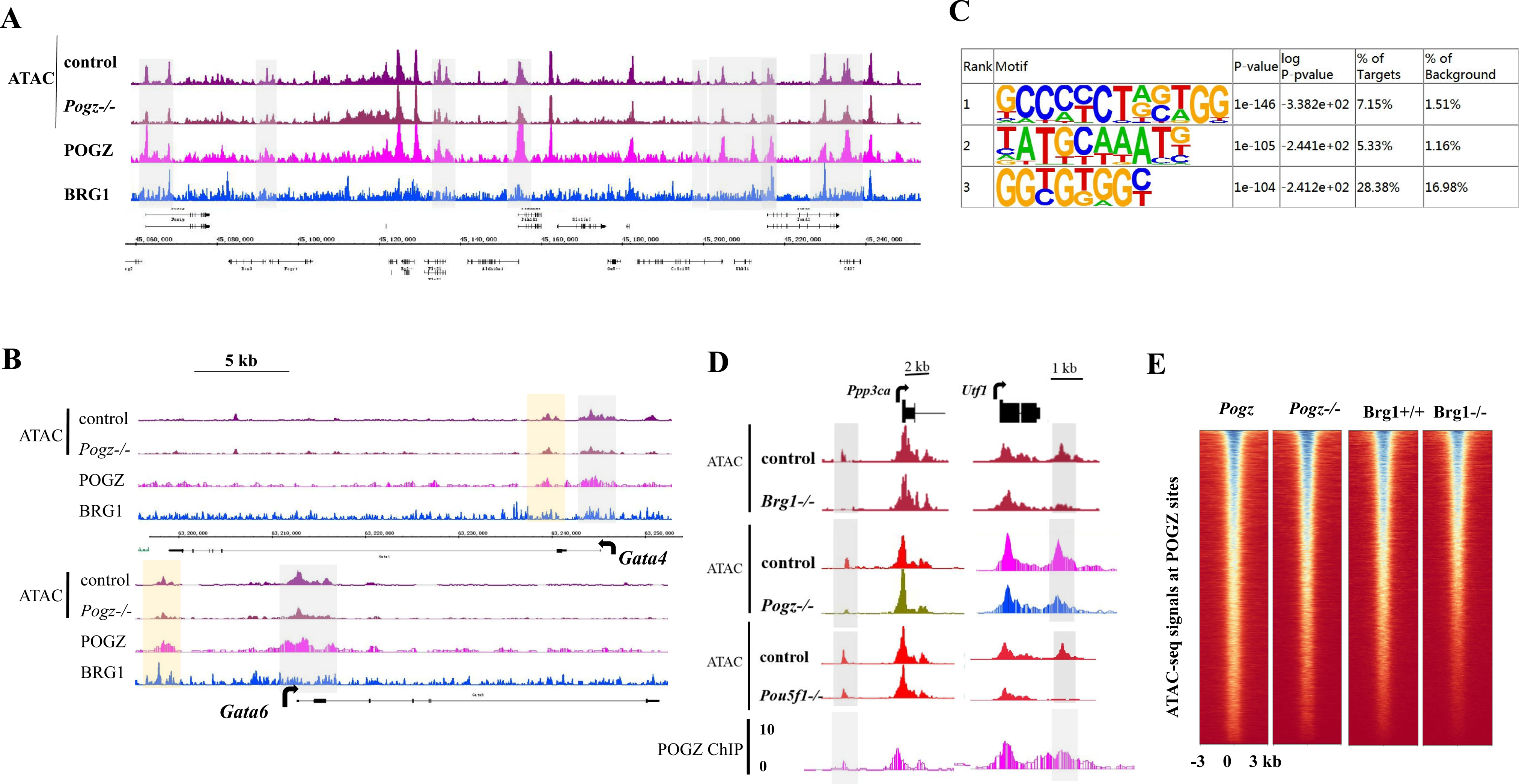
Related to Figure 7. **A** Snapshots of a randomly selected genomic region in chromosome 18 showing that the chromatin accessibility was reduced in the absence of POGZ at POGZ and BRG1 co-occupied sites. **B** Snapshots of the indicated gene loci showing that the chromatin accessibility was reduced in the absence of POGZ at POGZ and BRG1 co-occupied sites. Grey color highlighting the proximal TSS, and orange for distal enhancers. **C** The top three enriched motifs at sites where ATAC-seq signals were altered in the absence of POGZ by HOMER. **D** Snapshots of POGZ C&T and ATAC-seq signals at the indicated gene loci, showing that chromatin accessibility at POGZ-bound loci was similarly reduced in the absence of POGZ, BRG1 and OCT4. **E** Heat map of ATAC-seq signals at POGZ targets in control and BRG1-depleted ESCs (King and Klose, 2017). Sites were ranked by loss of ATAC-seq signals at POGZ-bound sites (n=10250).

